# Polycomb repressive complex 2 coordinates with Sin3 histone deacetylase complex to epigenetically reprogram genome-wide expression of *effectors* and regulate pathogenicity in *Magnaporthe oryzae*

**DOI:** 10.1101/2021.03.21.436344

**Authors:** Zhongling Wu, Jiehua Qiu, Huanbin Shi, Chuyu Lin, Jiangnan Yue, Zhiquan Liu, Wei Xie, Yanjun Kou, Zeng Tao

## Abstract

The strict suppression and reprogramming of gene expression are necessary at different development stages and/or in response to environment stimuli in eukaryotes. In Rice *Magnaporthe oryzae* pathosystem, *effectors* from pathogen are kept transcriptionally silenced in the vegetative growth stage and are highly expressed during invasive growth stage to adapt to the host environment. However, the mechanism of how such *effectors* are stably repressed in the vegetative stage and its roles during rice blast infection remain unclear so far. Here, we showed that all subunits of Polycomb Repressive Complex 2 are required for such repression by direct H3K27me3 occupancy and pathogenic process in *M. oryzae*. Suppression of polycomb-mediated H3K27me3 causes an improper induction of *effectors* during vegetative growth thus simulating a host environment. Notably, the addition subunit P55 not only acts as the bridge to connect with core subunits to form a complex in *M. oryzae*, but also recruits Sin3 histone deacetylase complex to prompt H3K27me3 occupancy for stable maintenance of transcriptional silencing of the target genes in the absence of PRC1. In contrast, during invasive growth stage, the repressed state of *effectors* chromatin can be partially erased during pathogenic development resulting in transcriptional activation of effectors therein. Overall, Polycomb repressive complex 2 coordinates with Sin3 histone deacetylase complex to epigenetically reprogram genome-wide expression of *effectors,* which act as molecular switch to memorize the host environment from vegetative to invasive growth, thus contributing to the infection of rice blast.

## Introduction

Organisms need to reprogram gene expression properly in different development stages or environment stimuli, which suppressing unnecessary genes is the most of important process (Cavalli and Heard, 2019, Netea et al., 2016). Stable maintenance of repressed gene expression states is achieved partly by the propagation of specific chromatin modifications (Cavalli and Heard, 2019). Evolutionary conserved polycomb repressive complexes (PRC) plays a vital role in such repression and contributes to facultative heterochromatin (Schuettengruber et al., 2017, Wiles and Selker, 2017, Margueron and Reinberg, 2011). PRC is involved in various biological processes, including maintaining cellular and tissue identity in multicellular organisms and regulating phase transitions in plants (Margueron and Reinberg, 2011, Blackledge et al., 2015). In mammals and higher plants, there are two main polycomb group complexes, PRC1 and PRC2 (Lanzuolo and Orlando, 2012). PRC1, including Pc, Ph, and Psc, compacts chromatin and catalyzes the monoubiquitylation of histone H2A. PRC2, including core subunits proteins, Ezh (Kmt6), Su(z)12, Esc (Eed) and additional subunits RbAp48/Nurf55 (P55), catalyzes the methylation of histone H3 at lysine 27 (Wiles and Selker, 2017, Schuettengruber et al., 2017). The core PRC2 complex is conserved from *Drosophila* to mammals and higher plants, while the PRC1 complex is not evolutionarily conserved. Notably, PRC1 has not been identified in fungi so far (Margueron and Reinberg, 2011, Ridenour et al., 2020).

In the fungi kingdom, disruption of PRC usually reprograms genome-wide gene expression, leading to abnormal growth and reduced pathogenicity (Ridenour et al., 2020, Wiles and Selker, 2017). In *Fusarium graminearum*, deletion of core subunits of PRC2 resulted in the complete loss of H3K27me3 modification, ∼2,500 genes up-regulation, as well as severe defects in growth and pathogenicity (Connolly et al., 2013). In *Magnaporthe oryzae*, deletion of *KMT6* eliminated all H3K27me3 modifications, resulted in a decrease in sporulation and highly reduced pathogenicity in wheat and barley (Pham et al., 2015). In *Neurospora crassa*, loss of core subunits also abolished all H3K27me3 modification, but only accompanied by slight growth defects (Jamieson et al., 2013). In the yeast *Cryptococcus neoformans*, loss of *EZH2, Eed1* or additional subunit *Bnd1* removed all H3K27me3 modification, while loss of the novel accessory component *Ccc1* resulted in regional reduction and relocalization of H3K27me3 modification on the chromosome (Dumesic et al., 2015). In symbiotic fungus *Epichloe festucae*, H3K27me3 coupled with H3K9me3 controls expression of symbiosis-specific genes, such as genes related to alkaloid bioprotective metabolites (Chujo and Scott, 2014). Although many studies have been done on the PRC complex in fungi, how the PRC2 effectively regulates gene expression and stably maintains the repressive H3K27me3 modification on the target chromatin in the absence of PRC1 remains largely unclear (Ridenour et al., 2020, Wiles and Selker, 2017).

During plant–pathogen interaction, pathogens have evolved multiple layer of strategies to invade host effectively, including secreting effectors to modulate rice immunity and ensure progressive infection (Fouche et al., 2018). Reprogramming the expression of the *effectors* by the pathogen is often used as an effective strategy to invade the host (Fouche et al., 2018). In addition, recent studies showed that transcriptional polymorphism with *effectors* contributes to the adaption of pathogenic microbes to environmental changes and co-evolution with host (Fouche et al., 2018). In the soybean root rot pathogen *Phytophthora sojae*, occupancy of H3K27me3 at *Avirulence* gene *Avr1b* leads to transcriptional silencing and fails to induce the *Rps1b* mediated disease resistance (Wang et al., 2020). However, in the vegetative growth stage, how to maintain the stable suppression of expression of *effectors* remains unclear.

Rice blast, caused by *M. oryzae*, is the most devastating disease of rice and threatens food security worldwide. In *M. oryzae*–rice interaction, the majority of the *effectors* are kept transcriptional silencing or expressed relatively low in the vegetative growth stage (Dong et al., 2015, Sharpee et al., 2017, Jeon et al., 2020), but are de-repressed during invasive growth stage (Dong et al., 2015, Jeon et al., 2020). During preparing our manuscript, a really good study found that H3K27 dynamics associate with altered transcription of *in planta* induced genes in *M. oryzae* (Zhang et al., 2021). Here, we not only find that histone modification H3K27me3 is required for genome-wide transcriptional silencing of *effectors* during vegetative growth stage in *M. oryzae*, but also present the detailed and novel mechanism beyond these findings. First, all subunits of PRC2, are required for H3K27me3-meditating polycomb silencing on the *effectors* and pathogenicity, which the addition subunit P55 acts as the bridge to connect with the core subunits to form a complex in *M. oryzae*. Furthermore, P55 recruits Sin3 histone deacetylase complex to coordinate with PRC2 and prompts H3K27me3 occupancy on the target chromatin. During invasive growth stage, the repressed chromatin state on the *effectors* can be partially “erased”, resulting in the transcriptional activation of *effectors*. In addition, we also reveal the important role of Sin3 histone deacetylase complex in transcriptional reprogramming the genome-wide expression of *effectors*, as well as in pathogenicity.

## Results

### Kmt6, as well as other PRC2 subunits, is indispensable for H3K27me3 modification in *M. oryzae*

In *M. oryzae*, Kmt6 (*MGG_00152*) was reported as the writer of H3K27me3 modification which is associated with altered transcription of *in planta* induced genes in *M. oryzae* (Pham et al., 2015, Zhang et al., 2021). However, how H3K27 trimethylation achieve stably transcriptional silencing of *effectors* during vegetative growth remains largely unknown. To answer this question, Δ*kmt6* was created in the wild-type (WT) *M. oryzae* strain B157. As reported, the expression levels of representative *effectors* during vegetative growth stage exhibited significant increased expression levels than that of the WT (Figure 1a)(Khang et al., 2010, Mosquera et al., 2009, Sharpee et al., 2017). To verify whether the increased transcriptional expression of *effectors* would enhance accumulation and secretion of effectors during vegetative growth stage, the *BAS4-GFP* construct was introduced into the Δ*kmt6* strain. As shown in the Figure 1b, the Bas4-GFP signal was clearly evident in the conidia, and the apical and subapical regions of the hyphae of the Δ*kmt6* mutant, but was not detectable in those of the WT. These results indicated that removal of H3K27me3 modification leads to untimely expression of *BAS4* during vegetative growth stage similar with that found naturally during invasive growth stage (Figure 1b).

**Figure 1.**
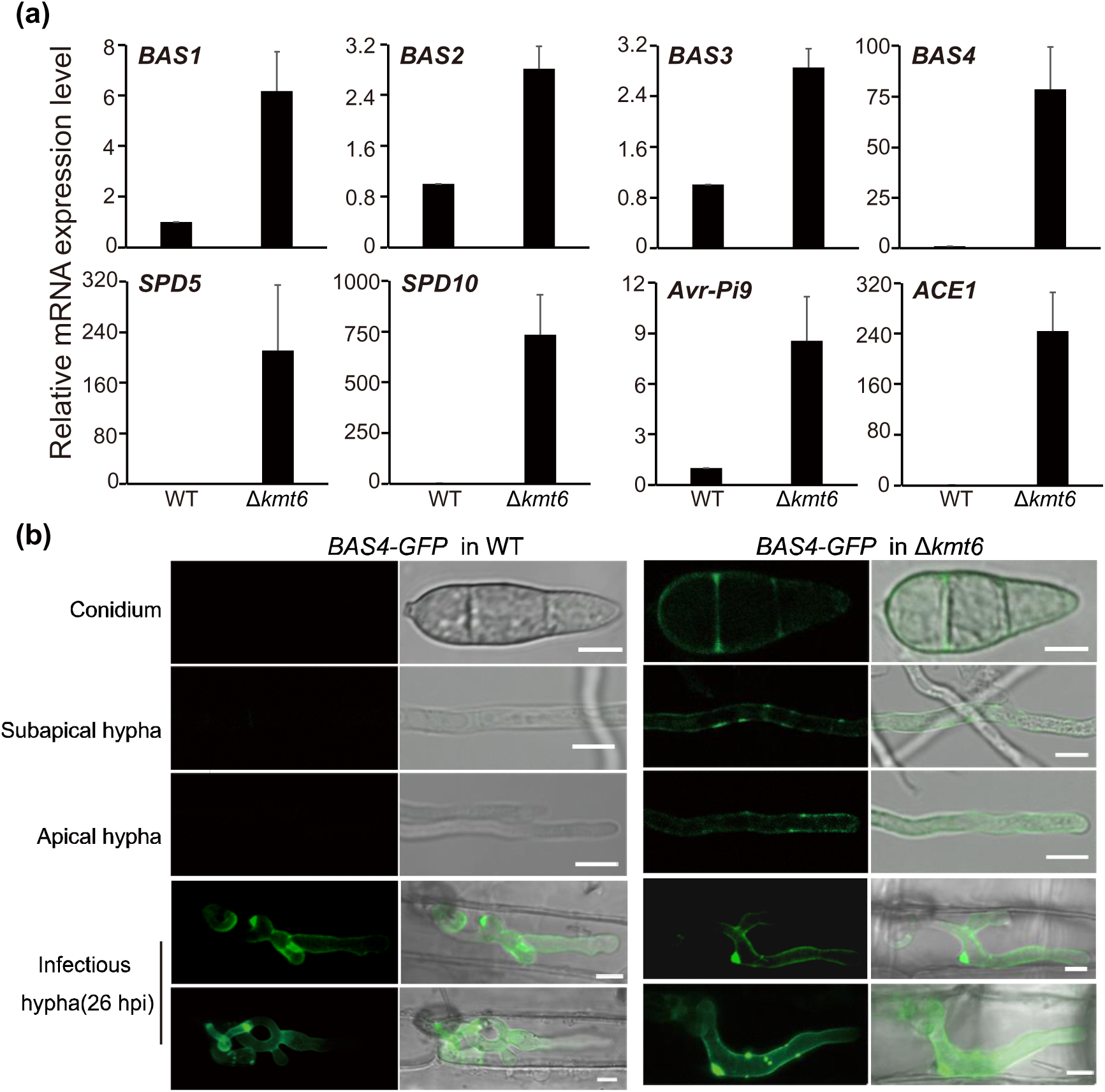
Loss of *KMT6* results in activation of *effectors* in the vegetative growth stage in *M. oryzae*. (a) Relative expression level of representative *effectors* in the WT and Δ*kmt6* strains by qRT-PCR. The *β-tubulin* gene was used as the endogenous reference gene. Bar means standard error of three biological repeats. (b) Confocal microscopy image of Bas4-GFP in the vegetative and *in planta* growth stages of the WT and Δ*kmt6* strains. In the WT, Bas4-GFP fluorescence is only presented in the *in planta* growth stage, while in the Δ*kmt6*, Bas4-GFP fluorescence could be detected in the conidia, apical hypha, subapical hypha and invasive hypha. Bar = 5 μm.

Polycomb-group genes were first identified in *Hox* regulation in *Drosohpila* and are usually assembled with Polycomb repressive complex 1(PRC1) and PRC2 (Blackledge et al., 2015, Ridenour et al., 2020). To identify the candidate PRC1 and other PRC2 subunits in *M. oryzae*, BLASTp was used to search for the orthologs from *M. oryzae* 70-15 genomes (taxid: 242507) with the query sequences from *Neurospora crassa, Arabidopsis thaliana,* and *Drosophila melanogaster* (Table S1). Three unique PRC2 core subunits, Kmt6 (MGG_00152), Eed (MGG_06028), Suz12 (MGG_03169), and one additional subunit P55 (MGG_07323) were obtained, while no PRC1 subunits was hit even by low stringency BLAST in *M. oryzae* (Table S1, Figure S1).

To investigate whether Kmt6, Eed, Suz12 and P55 form a complex, we first fused these proteins with GFP and examined their subcellular localization in *M. oryzae*. As shown in the Figure 2a, all these subunits were co-localized Hoechst-stained nuclei, suggesting that these components may form a complex in the nucleus like PRC2 in other species (Figure 2a). Then, yeast two-hybrid assay was conducted to test whether these subunits are physically associated *in vitro*. *KMT6*, *Suz12*, *Eed* and *P55* were cloned to the prey and bait vectors respectively. These results showed that the additional subunit P55 interacts with other three core subunits, and Kmt6 interacts with Eed in the yeast cells (Figure 2b). Furthermore, co-immunoprecipitation assays with strains expressing *Eed*-*GFP* and *Kmt6*-*Flag*, *P55*-*GFP* and *Kmt6*-*Flag*, *P55*-*GFP* and *Suz12*-*Flag*, *P55*-*GFP* and *Eed*-*Flag* respectively further confirmed these interactions *in vivo* (Figure 2c-f). Taken together, these results suggested that Kmt6, Eed, Suz12 and P55 form a complex, and P55 serves as the bridge to connect the PRC2 components.

**Figure 2.**
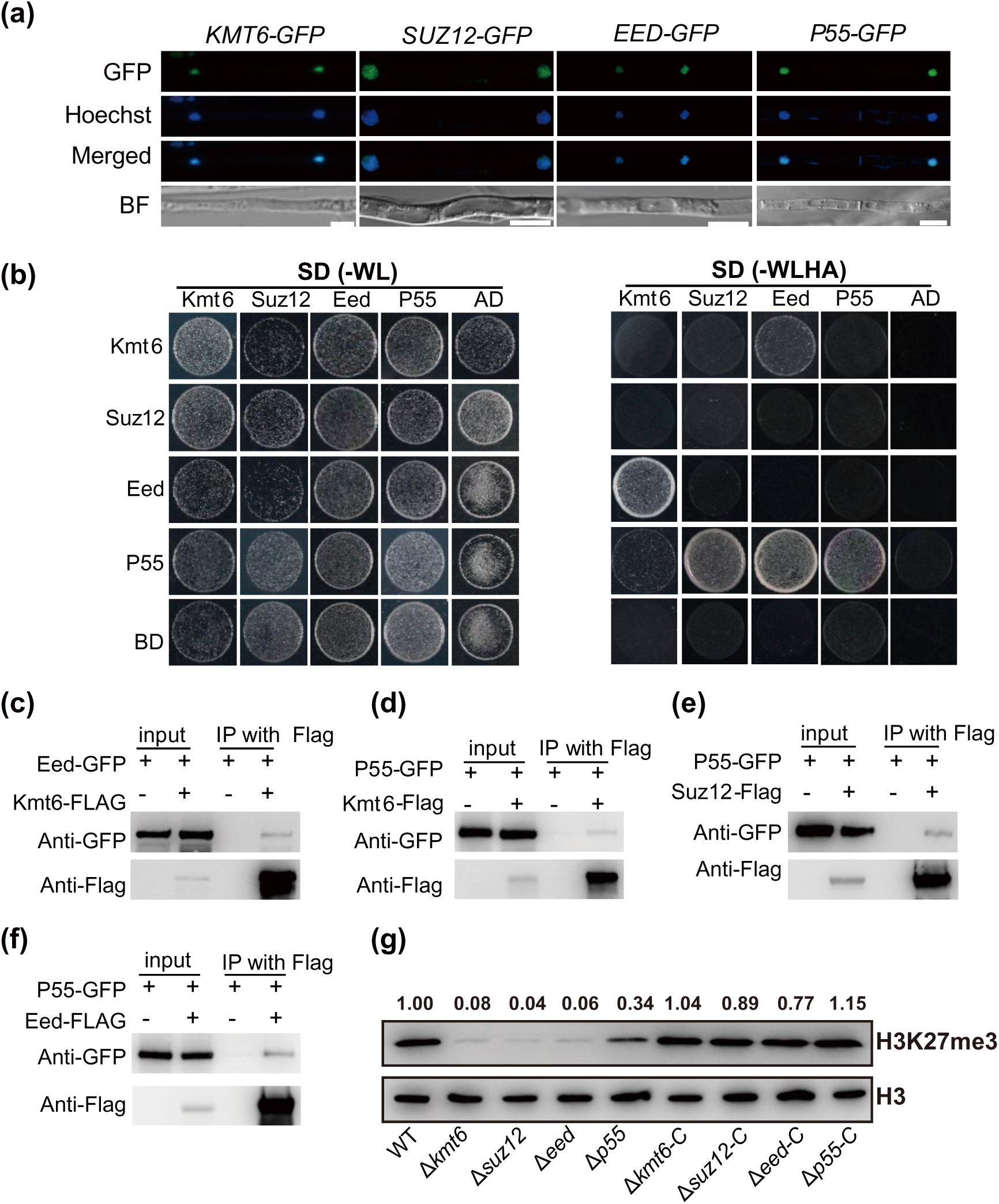
Core subunits Kmt6, Eed, Suz12, together with P55, are involved in the Poly-comb repressive complex 2 in *M. oryzae*. (a) Confocal microscopy-based subcellular localization of PRC2 subunits fused with GFP. The GFP signal co-localized with Hoechst (10 μg/mL) stained nuclei. Bar = 5 μm. (b) Yeast-two-hybrid assay of PRC2 components Kmt6, Eed, Suz12 and P55. The bait and prey plasmids were co-transformed into yeast strain Y2Hgold respectively, then the transformants were grown on basal medium SD (-WL, without tryptophan and leucine) and selective medium SD (-WLHA, without tryptophan, leucine, histidine and adenine). (c-f) Co-immuoprecipitation (CoIP) was performed with the strains expressing both *Kmt6-Flag* and *Suz12-GFP*, *P55-GFP* and *Kmt6-Flag, P55-GFP* and *Suz12-Flag*, or *P55-GFP* and *Eed-Flag*. (g) Western blot detection of histone H3 and histone methylation H3K27me3 in the WT, Δ*kmt6*, Δ*eed*, Δ*suz12* and Δ*P55* strains. The intensity abundance was measured and calculated relative to that of the WT. Two repeated biological experiments were carried out and the results were similar.

To explore whether Eed, Suz12, and P55 are indeed required for H3K27me3 modification, deletion mutants of *Suz12*, *Eed* and *P55* were created by homolog recombination in the WT. Subsequently, the protein levels of histone lysine methylation were detected with specific antibodies using Western blotting assay. The levels of H3K27me3 were almost undetectable in the Δ*kmt6*, Δ*eed* and Δ*suz12* mutants, while the level of H3K27me3 in the Δ*P55* still retained half level of the WT (Figure 2f), which indicated that P55 is not essential for H3K27me3 modification as the core subunits. Furthermore, the decreased levels of H3K27me3 in the mutants were completely restored in the complementary strains respectively (Figure 2f). Meanwhile the levels of H3K4me3 and H3K36me3 had no obvious change in all aforementioned strains by comparing with that of WT (Figure S2). We conclude that Kmt6, Suz12 and Eed are fully and specifically required for H3K27me3 modification, while P55 partially contributes to H3K27me3 modification.

### All PRC2 subunits are required for mycelia growth, conidiation, and pathogenicity in *M. oryzae*

To investigate the function of PRC2 complex in *M. oryzae*, mycelia growth and conidiation were assessed in the Δ*kmt6*, Δ*eed*, Δ*suz12*, Δ*p55*, as well as their complementary strains. Similar with the Δ*kmt6*, Δ*suz12* and Δ*eed* strains exhibited decreased mycelia growth and dramatically reduced conidiation, which had similar phenotype with recent report (Zhang et al., 2021) (Figure 3a-c). Notably, Δ*p55* had more severe phenotype in the mycelia growth (Figure 3a-b), suggesting that P55 had wider functions beside for acting as the bridge in the PRC2 complex. These results indicated that H3K27me3 is required for the mycelia growth and conidiation in *M. oryzae*.

**Figure 3.**
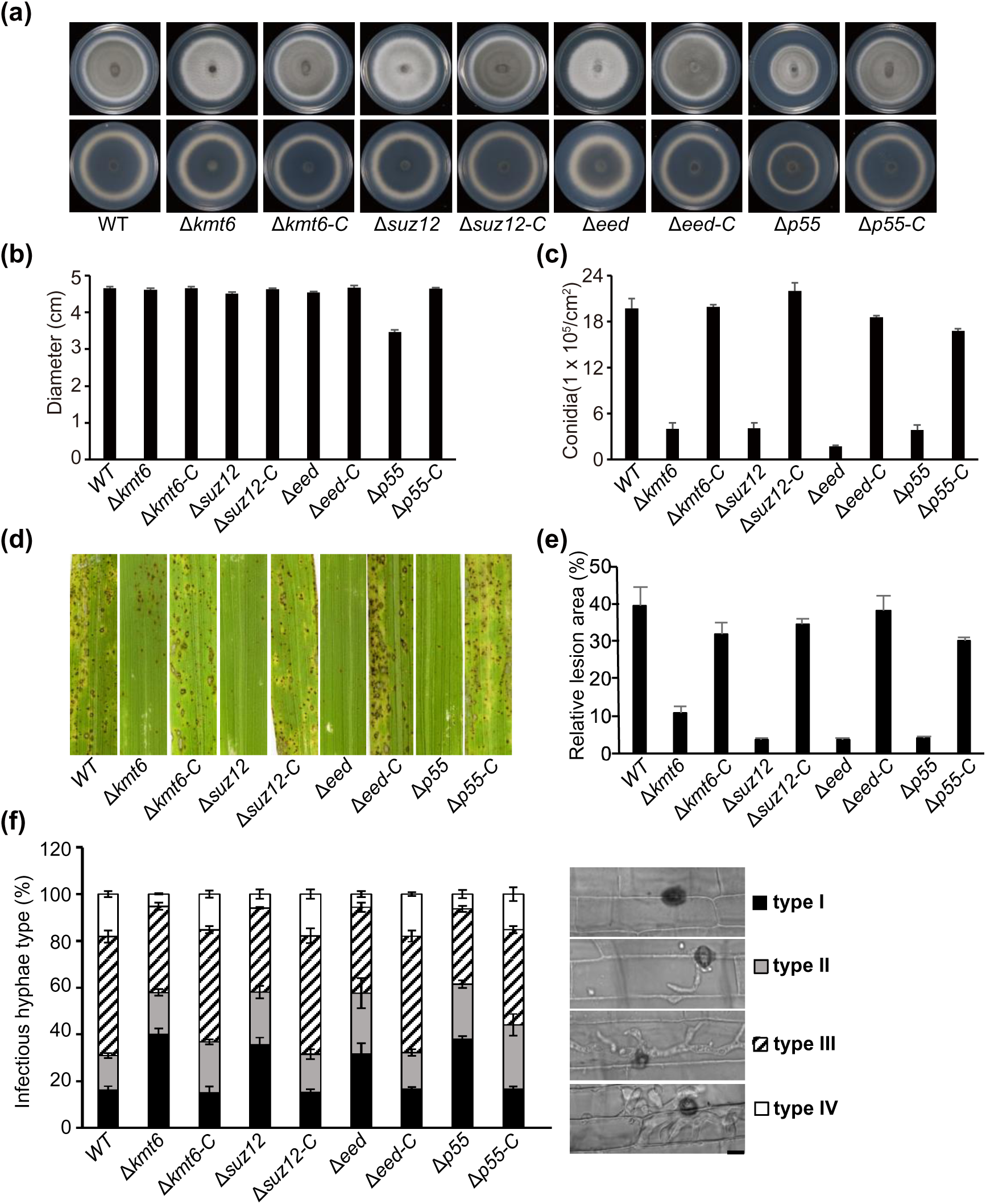
PRC2 complex is required for fungal growth and pathogenicity in *M. oryzae*. (a) Radical growth of the WT, Δ*kmt6*, Δ*eed*, Δ*suz12*, Δ*P55* and their complementation strains. Colonies of indicated strains were grown on the PA medium for 7 d, then both up and bottom sides of colonies were imaged. (b) Statistical analysis of colonies diameters of tested strains on the PA medium. (c) Bar charts showing the conidia that produced in the indicated strain of *M. oryzae*. (d) Observation and statistical analysis of invasive hypha growth in rice sheath cells at 40 hpi. Four types of invasive hypha (illustrated in the right panel with corresponding column): no penetration, penetration with primary hyphae, with differentiated secondary invasive hyphae, and invasive hyphae spreading into neighbouring cells, were quantified. Data represents mean ± SD of three independent repeats, with n = 300 appressoria per analysis. Scale bar = 5 μm. (e) Blast infection assays using rice seedlings (*Oryza sativa* L., cultivar CO39). Deletion of *KMT6*, *EED*, *SUZ12* and *P55* impaired the pathogenicity of *M. oryzae*. (f) Relative lesions area of indicated strains. The relative area of lesions was quantified by Image J software. Data represents the mean ± SD from three independent replicates.

To analyse the function of PRC2 components in pathogenicity, conidia of the WT, mutant and complementary strains were collected and then inoculated on the susceptible rice cultivar CO39 (*Oryza sativa*). As the number of conidia from mutants highly reduced (15-25% of the WT) (Figure 3c), low concentration of conidia (5 × 10^4^/ mL) were used in the rice seedling infection assay. Compared with WT, which caused the characteristic spindle-shaped blast lesions with grey centres, mutant strains formed relative fewer and restricted lesions (Figure 3d-e). To further decipher these observations, the appressoria formation on the inductive hydrophobic surface and rice sheath were investigated. Although no obvious change in appressoria formation at 24 hpi (hour post inoculation) was found in the mutant strains (data not shown), the invasive growth displayed significant difference between the mutants and WT (Figure 3f). Nearly 90% appressoria from the WT strain successfully penetrated the rice sheath, while only 60% appressoria from the deletion mutants were capable of penetration (Figure 3f) at 40 hpi. Moreover, the formation of secondary invasive hyphae in the mutant strains was less than that of WT. These results indicated that H3K27me3 is necessary for penetration and invasive hyphal growth.

### Core subunits of PRC2, together with P55, suppress genome-wide gene expression, including *effectors*, at vegetative growth stage

To explore the roles of H3K27me3 in the transcriptional regulation in *M. oryzae*, high-throughput sequencing (RNA-seq) was performed using the vegetative mycelia with three biological repeats. As H3K27me3 always contributes to transcriptional silencing on target genes, the mutant strains would be expected to enrich with high ratio of de-repressed genes. As expected, totally 19.2%, 18.6%, 20.1% and 15.4% of genome-wide genes were identified as up-regulated genes (UEG) with equal or greater than 2-fold change (Log_2_ > 1, P<0.05), while only 5.3%, 6.0%, 8.8% and 3.3% of genes were identified as down-regulated genes (DEG) in the Δ*kmt6*, Δ*eed*, Δ*suz12* and Δ*p55* strains respectively (Log_2_ <-1, P<0.05) (Figure 4a). As the core subunits of PRC2, Δ*kmt6*, Δ*eed* and Δ*suz12* shared 80-86% overlaps (2144 genes) in their UEG (Figure S3a). The high ratio of co-regulated genes further indicated that Kmt6, Eed and Suz12 function in the same complex. Moreover, the predominant roles of de-repressed regulation in the mutants of core subunits suggested that H3K27me3 modification played conserved roles as transcriptional repressors in *M. oryzae* (Figure 4a). Although P55 was thought as additional subunit which was different from other three core subunits, there was no obvious difference in the gene sets between the Δ*kmt6*-UEG and Δ*p55*-UEG strains (P=0) (Figure S3b). These results further confirmed that core subunits of PRC2, together with P55, function in the same complex and regulate similar biological processes.

**Figure 4.**
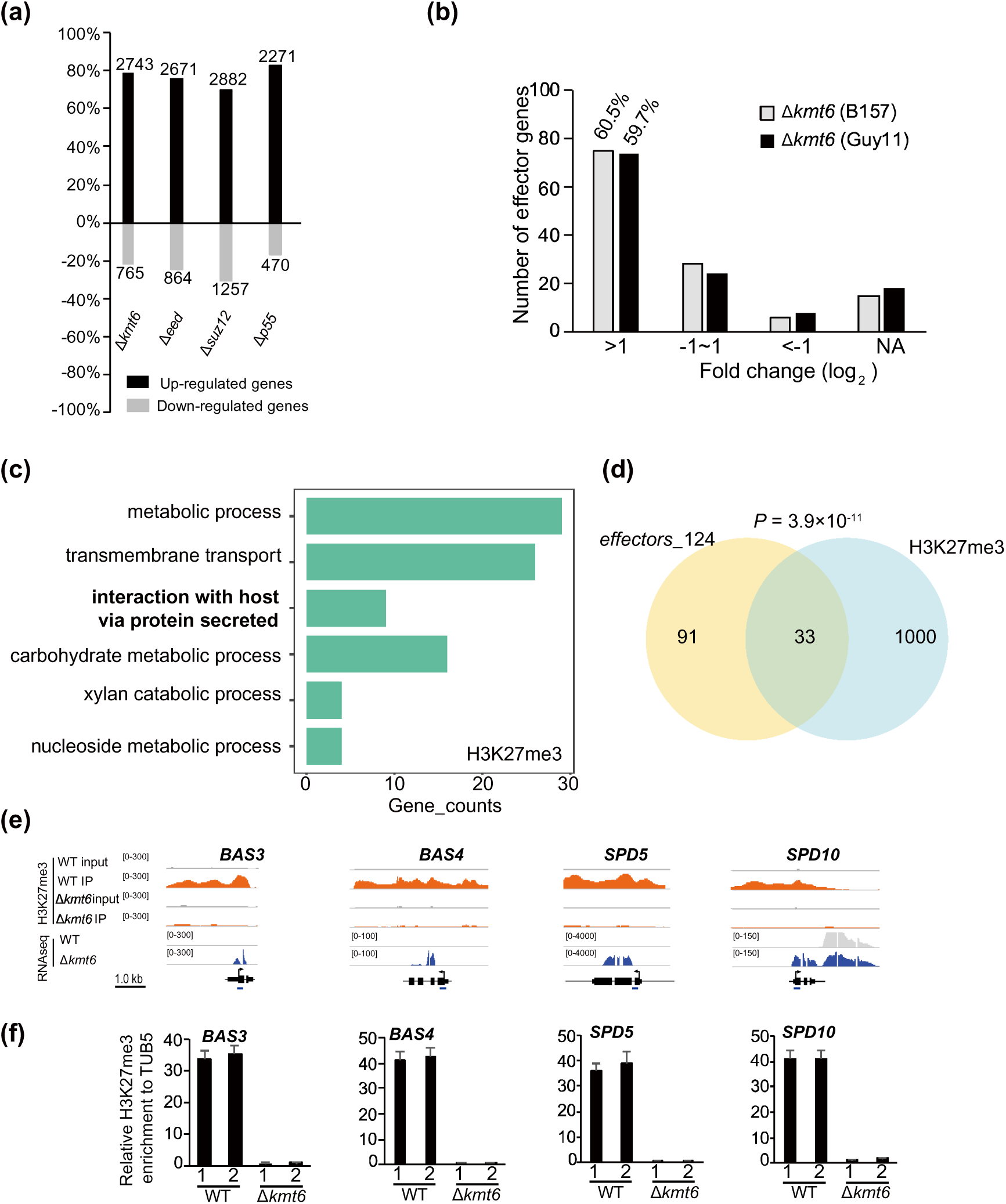
PRC2-mediated H3K27me3 activity is necessary for the suppression of *effectors* during vegetative growth stage in *M. oryzae*. (a) Summary of up- and down-regulated genes in the different *PRC2* component mutants with RNA-seq analysis. The number at the top or bottom is the number of DEGs in the mutant. The Y-axis is the percentage of up- and down-regulated genes in all DEGs. Data was obtained from three independent biological repeats. (b) The distribution of 124 *effectors* based on the expression change in Δ*kmt6*-UEG (B157) and Δ*kmt6*-UEG (Guy11) compared with that of their WT respectively with our RNA-seq analysis. (c) GO analysis of H3K27me3 marked genes. (d) The Venn diagram showing statistically significant overlap between gene sets of H3K27me3 marked genes and putative 124 *effectors.* (e) Genome browser views of H3K27me3 occupancy from ChIP-seq and expression pattern from RNA-seq of selected genes. ChIP-seq was conducted with two biological repeats. The number areas were reads per million (RPM). (f) ChIP-qPCR verified the enrichment of H3K27me3 at the chromatin of selected *effectors*. The examined regions were shown at the bottom. The relative enrichments were calculated by relative quantitation from two biological repeats, which was standardized with internal control *TUB5*, and then compared with that of WT. Bar represents standard error from three technical repeats.

To explore the detailed biological process involved in PRC2, Gene Ontology (GO) analysis was conducted with Δ*kmt6*-UEG. Biological process as “interaction with host *via* secreted protein” were significantly enriched (Figure S3c). This was consistent with our hypothesis that H3K27me3 modification contributes to suppress the expression of *effectors* at vegetative growth stage. To further address this question, gene sets of putative genome-wide *effectors*, including 134 (only 124 *effectors* had reads in our RNA-seq) and 247 (only 246 *effectors* had reads in our RNA-seq) *effectors* from two different literatures respectively, were extracted for further analysis (Sharpee et al., 2017, Dong et al., 2015). Compared with the WT, nearly 51.6% (64/124) and 42.7% (105/246) *effectors* exhibited significantly increased expression in the Δ*kmt6* strain (Figure 4b and S4). To preclude the specificity of strain background, similar analysis was performed with the Δ*kmt6* mutant in the Guy11 background. The ratio of up-regulated *effectors* in the Δ*kmt6* either in B157 or Guy11 background were significantly enriched with similar tendency (Figure 4b and S5a-c). We concluded that H3K27me3 is specifically required in the suppression of *effectors* during vegetative growth stage in *M. oryzae*.

In addition, loss of H3K27me3 modification not only activated the expression of *effectors*, but also activated some infection-specific genes such as those encoding cutinases and cell-wall degrade enzymes, which indicated that the absence of H3K27me3 could partially mimic host-derived signal. Therefore, we compared the gene sets between Δ*kmt6*-UEG and UEG from 36 hpi and 72 hpi *in planta* from published RNA-seq respectively (Jeon et al., 2020). The results showed that 30.1% (820/2654) of 36 hpi-UEG and 33.1% (778/2353) 72 hpi-UEG were significantly overlapped with Δ*kmt6-*UEG (Figure S6a-b).

### H3K27me3 occupancy is associated with increased transcription in the PRC2 mutants

To further depict whether H3K27me3 directly deposits on the chromatin of *effectors*, we attempted to map the genome-wide H3K27me3 occupancy using chromatin immunoprecipitation assay followed by high-throughput sequencing (ChIP-seq) in the WT and Δ*kmt6* strains. The intensity of H3K27me3 occupancy in the Δ*kmt6* mutant was barely detectable (Figure S7a), which was consistent with the role of Kmt6 as the H3K27me3 writer in *M. oryzae.* Compared with the Δ*kmt6* mutant, totally 1082 significant peaks were identified in the WT (Log_2_ > 1, P < 0.05), which were corresponded to 1033 genes and mainly spread within gene bodies (Supplemental Figure 7a-b). To verify whether Δ*kmt6-*UEG were directly associated with the loss of H3K27me3 occupancy, the gene sets between Δ*kmt6-*UEG under low or high threshold and H3K27me3-marked genes were compared. With low threshold (Log_2_ > 1, P < 0.05), 23.5% (645 of 2743) of Δ*kmt6-*UEG were marked with directly H3K27me3 occupancy (Figure S7c). while with stringent threshold (Log_2_ > 3, P < 0.05), 40.1% (532 of 1327) of Δ*kmt6-*UEG were marked with H3K27me3 (Figure S7c). Together, the de-repressed genes in the Δ*kmt6* are highly correlated with the absence of H3K27me3 occupancy.

Next, GO analysis was conducted with H3K27me3-occupancy genes. Biological process “interaction with host via protein secreted” were highly enriched (Figure 4c), similar with the results from the Δ*kmt6-*UEG. Significantly, 26.7% (33 of 124) and 19.9% (49 of 246) of effectors were directly marked with H3K27me3 modification (Figure 4d and S7d). The chromatin of representative *effectors* as *BAS3*, *BAS4*, *SPD5* and *SPD10* enriched with H3K27me3 modification in the WT but nearly undetectable in the Δ*kmt6* mutant (Figure 4e). Consistently, the increased expression of examined *effectors* accompanied with almost undetectable H3K27me3 level in the Δ*kmt6* with ChIP-qPCR assay (Figure 1a and 4f). In addition, gene sets of 36 hpi-UEG and 72 hpi-UEG *in planta* were also highly marked with H3K27me3 (Figure S6a-b) and the chromatin state of four cutinases loci was validated with ChIP-qPCR (Figure S7e). These results further indicated that the H3K27me3 modification directly associates with transcriptional silencing on the *effectors* and infection-specific genes.

### P55 recruits Sin3 histone deacetylase complex to form a co-suppression complex

How PRC2 performs stable transcriptional silencing on the target chromatin in the absence of PRC1 remains unclear (Ridenour et al., 2020, Margueron and Reinberg, 2011). As a chromatin assembling factor and histone-binding WD40-repeat proteins, ortholog of the additional subunit P55 was reported to associate with other co-suppression complex to conduct transcriptional silencing in other organisms (Mehdi et al., 2016, Gu et al., 2011). As the additional subunit of PRC2 in *M. oryzae*, P*55* acts as the bridge to connect other core subunits together and Δ*p55* exhibited more severe phenotype than other mutants in our experiments. We postulated that P55 may recruit other co-suppression machinery to help PRC2 to stably maintain polycomb silencing. To address this possibility, P55 was fused to the bait vector and putative histone deacetylases were cloned individually in the prey vector for yeast-two-hybrid assays. As a result, we found that P55 interacted with four components of Sin3 histone deacetylases complex in yeast two-hybrid assays (Figure 5a). Sin3 histone deacetylases complex has been well characterized in yeast and other organisms (Huang et al., 2019, Grzenda et al., 2009, Adams et al., 2018), which the core components of Sin3 histone deacetylase complex includes histone deacetylase Hos2 (MGG_01633)(Lee et al., 2019), major regulatory protein Sin3 (MGG_13498), other subunits Sap18 (MGG_05680) and Sap30 (MGG_11142) in *M. oryzae*. Moreover, Sap30 interacted with other three components as Hos2, Sin3 and Sap18 in yeast which indicated that these components function in the same complex (Figure S8a). Coimmunoprecipitation assays with the strains expressing *P55*-*GFP*/*Sin3*-*Flag*, and *P55*-*GFP*/*SAP18*-*Flag* further confirmed that P55 is physically associated with Sin3-HDAC *in vivo* (Figure 5b-c).

**Figure 5.**
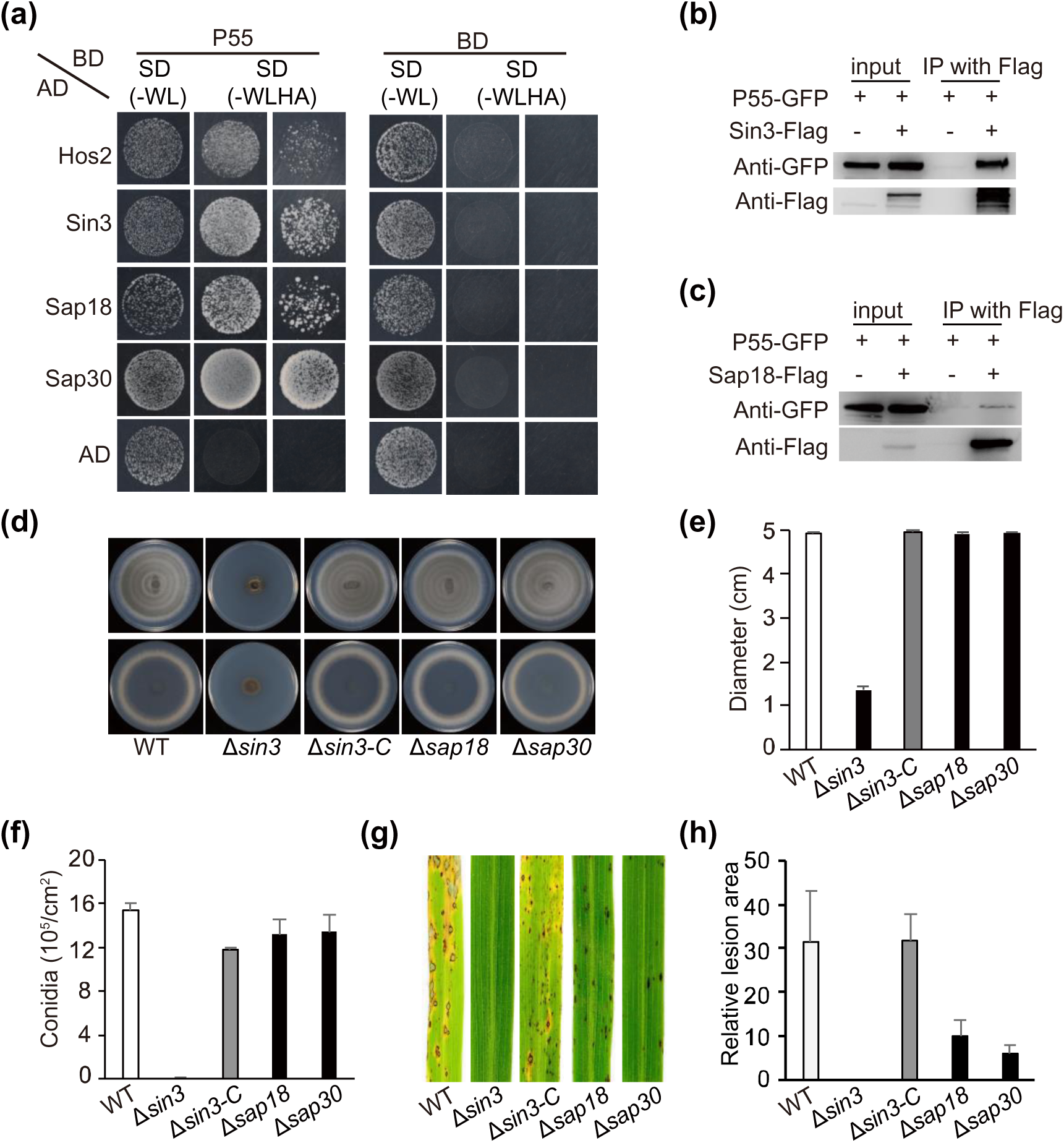
P55 associates with Sin3-HDAC complex which is required for pathogenicity in *M. oryzae*. (a) Yeast-two-hybrid assay of P55 and Sin3-HDAC component HDAC, Sin3, Sap18, and Sap30. The bait and prey plasmids were co-transformed into yeast strain Y2Hgold respectively, then the transformants were grown on basal medium SD (-WL) and selective medium SD (-WLHA). (b-c) Co-immunoprecipitation assays were performed with the strains expressing Sin3-Flag and P55-GFP, Sap18-Flag and P55-GFP. (d-e) Radical growth of the *SIN3-HDAC* deletion strains on the PA medium cultured for 7 d. (f) Deletion of *Sin3* resulted in decreased conidiation. (g-h) *Sin3-HDAC* is required for pathogenicity of *M. oryzae*. Rice seedling infection assay of the WT and *Sin3-HDAC* deletion strains with cultivar Co39. Similar results were obtained from two biological repeats.

To explore the function of Sin3-HDAC in *M. oryzae*, the deletion mutants of *Sin3*, *Sap18* and *Sap30* were created. Among these mutants, Δ*sin3* exhibited severe phenotype similar to Δ*p55*, including decreased fungal growth, severely reduced conidiation, as well as highly reduced pathogenicity (Figure 5d-h). To investigate whether Sin3-HDAC contributes to the histone deacetylation activity *in vivo*, the global levels of histone acetylation were checked in the WT and Δ*sin3* strains. Among the examined acetylated residues, at least the levels of H3K4ac, H3K27ac and H4K5ac were significantly increased in the Δ*sin3* strain than that of WT (Figure 6a and S8b). The increased level of H3K4ac in the Δ*sin3* was further confirmed by ChIP-seq assay. The global H3K4ac occupancy in the Δ*sin3* was significantly increased than that of WT, which further confirmed that Sin3-HDAC has histone deacetylation activity in *M. oryzae* (Figure S8c). As loss of histone deacetylation would accompany with genome-wide gene activation, we conducted RNA-seq with total RNA extracted from the Δ*sin3* and WT strains. In the vegetative growth stage, deletion of *Sin3* dramatically altered the expression of nearly 38% of genome-wide genes compared with WT. Totally, 3049 genes were up-regulated (Log_2_ > 1, P < 0.05), and 1454 genes were down-regulated (Log_2_ < −1, P < 0.05) which implied that *Sin3* also acts as a transcriptional repressor similar with P55 and Kmt6 (Figure 6b). Based on these results, we conclude that Sin3-HDAC contributes to histone deacetylation activity and acts as a transcriptional repressor to regulate genome-wide gene expression in *M. oryzae*.

**Figure 6.**
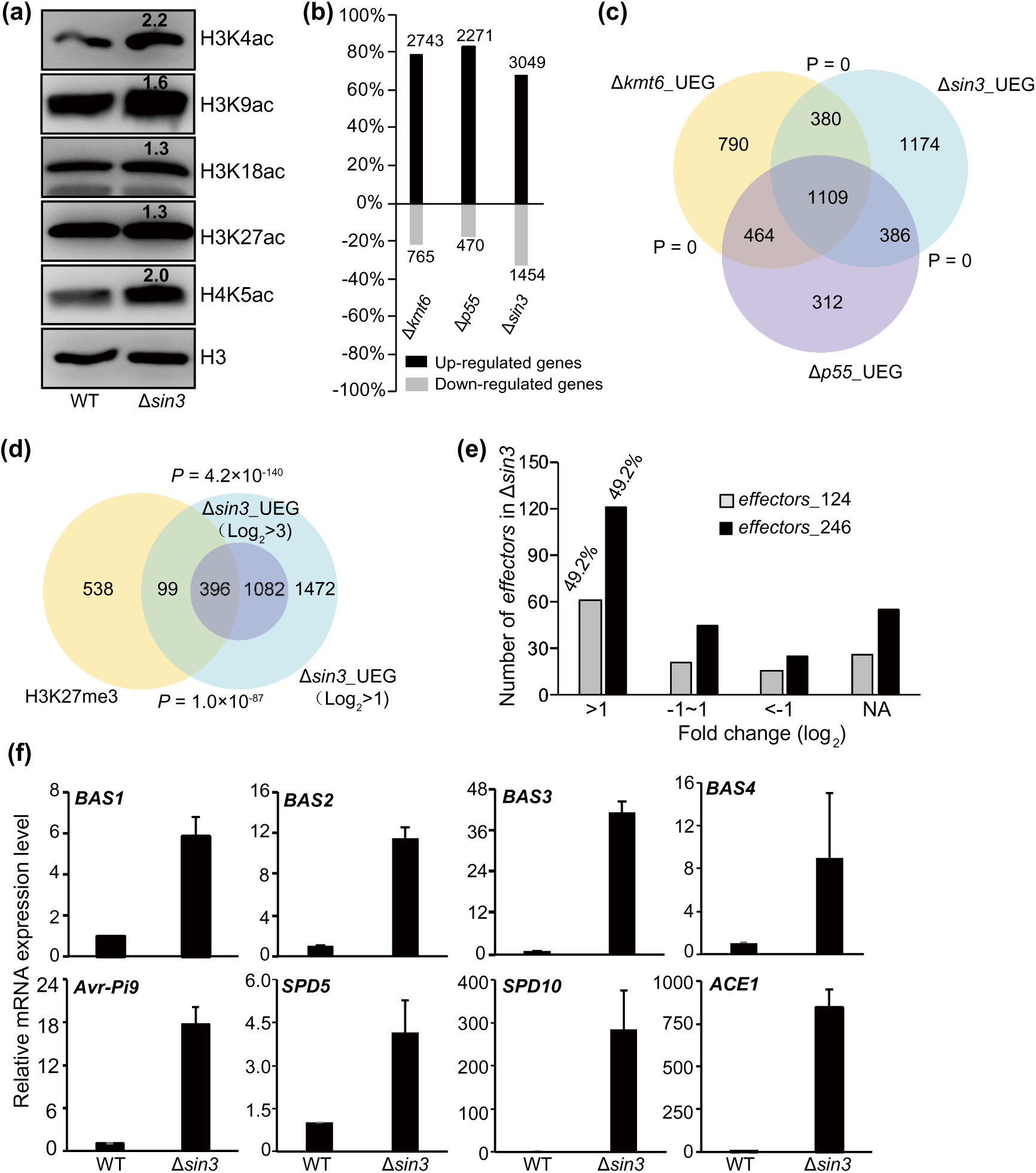
Sin3-HDAC coordinates with PRC2 to prompt H3K27me3-mediated transcriptional silencing on *effectors*. (a) Levels of histone acetylation at the examined residues were measured by Western blotting assay in the WT and Δ*sin3* strains. The number at the top of band is the relative intensity calculated by Image J software. Similar results were obtained from three biological repeats. (b) Summary of up- and down-regulated genes in the Δ*p55* and Δ*sin3* by comparing with that of WT. The number at the top or bottom is the number of DEG in the mutant. The Y-axis is the percentage of up- and down-regulated genes in all DEG. Data was obtained from three biological repeats. (c) The Venn diagram showing statistically significant overlaps among genes sets of Δ*sin3*-UEG, Δ*p55*-UEG and Δ*kmt6*-UEG. P value with Fisher’s exact test for overlapping between gene sets were labeled. (d) The Venn diagram showing statistically significant overlap between genes sets of H3K27me3-marked genes and Δ*sin3*-UEG. (e) The distribution of putative 124 and 246 *effectors* based on the expression change in Δ*sin3*-UEG. (f) Relative expression level of representative *effectors* were checked in WT and Δ*sin3* by qRT-PCR. Data represents mean±SD of three independent biological replicates.

### Sin3-HDAC co-operates with PRC2 to prompt transcriptional repression on the target genes

To explore the transcriptional correlation between Sin3-HDAC and PRC2, the transcriptional profiles of the Δ*sin3* and Δ*kmt6* strains were compared. A total of 1489 genes which were 54.3% of Δ*kmt6*-UEG and 48.8% of Δ*sin3*-UEG were significantly overlapped (P=0) (Figure 6c). To further address whether Sin3-HDAC contributes to H3K27me3-mediated transcriptional silencing, we investigated whether Δ*sin3*-UEG are enriched with H3K27me3 occupancy. The results showed that with lower stringent threshold (Log_2_ > 1, P < 0.05), 16.2% (495 of 3049) of Δ*sin3*-UEG were occupied with H3K27me3 modification. While with higher stringent threshold (Log_2_ > 3), 26.8% (396 of 1478) of Δ*sin3*-UEG were directly associated with H3K27me3 occupancy (Figure 6d). Meanwhile, enhanced H3K4ac and H4K5ac levels were also observed in the Δ*kmt6* and other PRC2 deletion mutants (Figure S8d). These results suggested that Sin3-mediated histone deacetylation contributes to PRC2-mediated transcriptional silencing on the target genes.

Next, GO analysis was conducted with Δ*sin3*-UEG. The term of “interaction with host via protein secreted” were highly enriched (Figure S9), which was similar with the results from RNA-seq in the Δ*kmt6* strain, indicating that Sin3-HDAC and PRC2 may coordinately regulate the expression *of effectors.* Therefore, the putative effectome that includes 124 and 246 predicted members were further investigated in the Δ*sin3*-UEG (Dong et al., 2015, Sharpee et al., 2017). The results showed that 49.2% (61 of 124, 121/246) predicted *effectors* exhibited significantly increased expression in the Δ*sin3* strain (Figure 6e). To verify these results from RNA-seq analysis, RT-qPCR was performed with three independent repeats in the WT and Δ*sin3* strains. Consistently, the expression of the *BASs* and *SPDs* in Δ*sin3* were significantly de-repressed in the vegetative growth stage (Figure 6f). In addition, a precocious and significant induction of *BAS4-GFP* signal was found in the conidia and vegetative hyphae of Δ*sin3*, thus indicating that deletion of *Sin3* leads to Bas4 accumulation and secretion during the vegetative growth stage (Figure S10).

Taken together, these results conclude that P55 recruits Sin3-HDAC as a co-repressor complex to maintain stably transcriptional silencing on the target genes, including a large scale of *effectors,* during vegetative growth in *M. oryzae*.

### H3K27me3 modification is “erased” from the chromatin of *effectors* loci during *M. oryzae*–rice interaction

To address whether the activated expression of *effector*s during infection stage is associated with reduced H3K27me3 occupancy, we attempted to conduct ChIP-qPCR to check H3K27me3 enrichment on chromatin of the *effectors* as *BAS3*, *BAS4*, *SPD5* and *SPD10* using *in vivo* infected rice leaves with the WT strain. The results showed that expression of four tested *effectors* was significantly induced from 24 hpi to 72 hpi (Figure 7a). Subsequently, 72 hpi-infection leaves were collected for ChIP assay. Compared with the mycelia, the relative H3K27me3 abundance of 72 hpi-infection leaves dramatically reduced at the chromatin regions of *BAS3*, *BAS4* and *SPD5*, except *SPD10*, which indicated that the activated expression of *effectors* would be partially caused by removing of H3K27me3 modification (Figure 7b).

**Figure 7.**
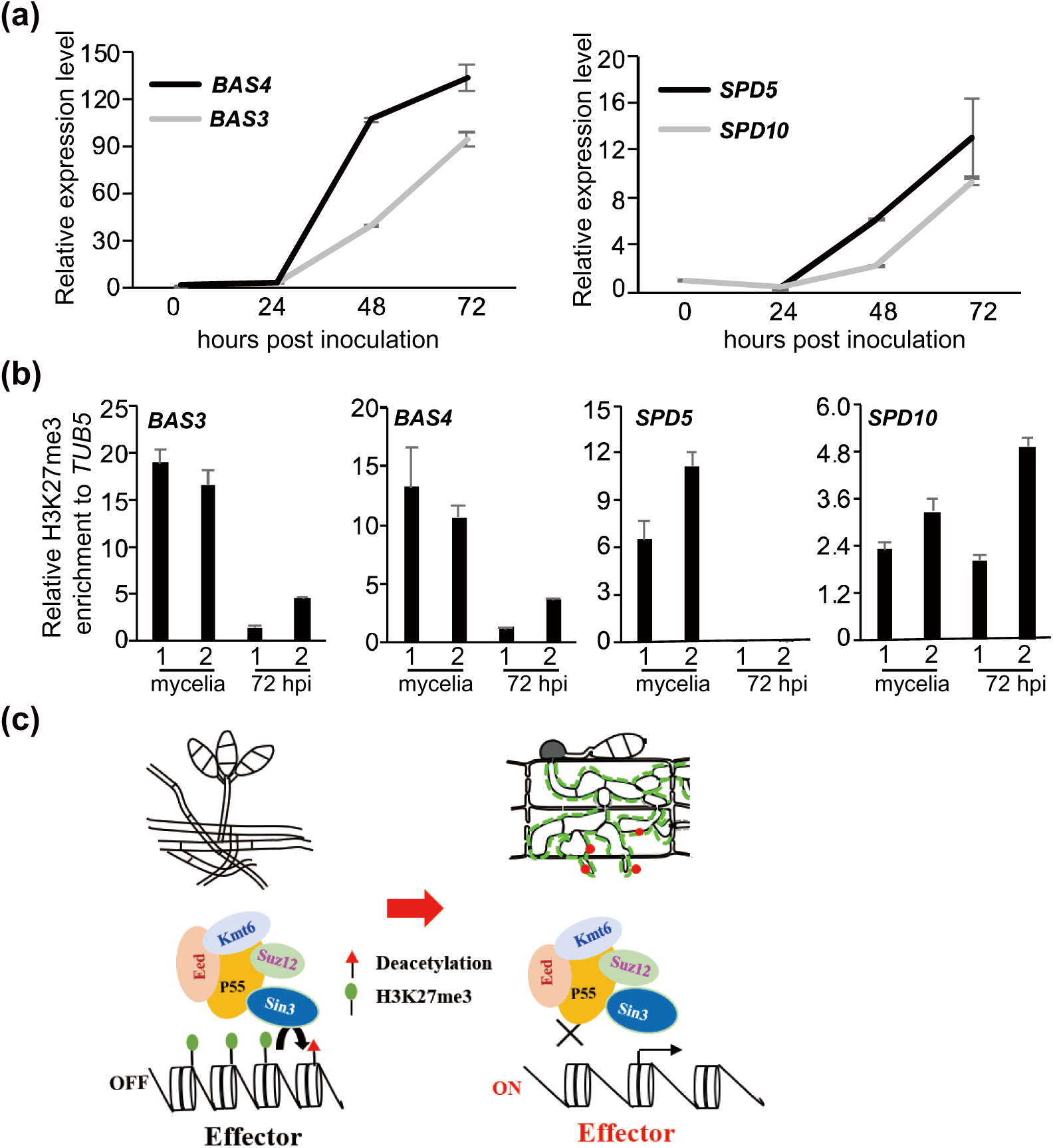
H3K27me3 modification is “erased” from the chromatin of *effectors* at the early stage of infection. (a) The expression of the *effectors BAS3*, *BAS4*, *SPD5*, and *SPD10* were induced by host. (b) H3K27me3 modification at the chromatin of *effectors* was removed at the early stage of infection. (c) Model of *effectors* reprogramming during transition from vegetative growth to *in planta* growth. During vegetative growth stage, PRC2 coordinates with Sin3 histone deacetylase complex suppresses the expression of *effectors* in *M. oryzae*, while during the invasion growth stage, H3K27me3 modification is “erased” upon the perception of signals from host, then the *effectors* are expressed.

## Discussion

In this study, we have revealed how H3K27me3 modifies the expression of genome-wide *effectors* and contributes to infection of *M. oryzae*. Suppression of H3K27me3 improperly activated the expression of *effectors* in the vegetative growth stage, as well as highly decreased pathogenicity. We also revealed a new mechanism that the additional subunit P55 acts as the bridge to connect core subunits to form a complex and recruits Sin3-HDAC to prompt H3K27me3 occupancy on the *effectors*, therefore stably maintain polycomb silencing in the absence of PRC1. During invasive growth stage, the repressed chromatin state on *effectors* is partially “erased”, subsequently the expression of *effectors* is activated to promote the infection of *M. oryzae*.

Epigenetic modification was correlated with genome plasticity and stability, help pathogen effective adaption to host environments (Chadha and Sharma, 2014, Wang et al., 2020, Cavalli and Heard, 2019). In *M. oryzae*, epigenetic modification as histone (de)methylation and (de)acetylation have been extensively explored in different biological processes such as infection structure morphogenesis, developmental transition and autophagy. Histone deacetylases Tig1 and Hos2 regulate infectious growth and asexual development in *M. oryzae* (Ding et al., 2010, Lee et al., 2019). H3K4 methyltransferase Set1 contributes to appressorium formation, conidiation and pathogenicity with transcriptional activation on the infection-related genes (Pham et al., 2015). Histone acetyltransferase Gcn5 negatively regulates light-induced autophagy and conidiation through acetylated the autophagy protein Atg7 (Zhang et al., 2017). *MoSNT2* plays key roles in infection-associated autophagy through recruiting histone deacetylase complex on the target chromatin (He et al., 2018). Recently, Zhang et al. elucidated that H3K27 dynamics associate with altered transcription of *in planta* induced genes, including *effectors*, in *M. oryzae* (Zhang et al., 2021). In our study, we found that all core subunits of PRC2, together with the additional subunit P55, are required for H3K27me3-meditating polycomb silencing on the *effectors* in the vegetative growth stage and pathogenicity. Our results also revealed that histone modification H3K27me3, coordinately mediated by PRC2 and Sin3 histone deacetylase complex, directly suppresses most of the genome-wide *effectors* expression. Previous studies suggested that pathogen *effectors* are rapidly evolved and especially distributed in regions with high genome plasticity. In *M. oryzae*, *effectors* do not have common motifs or conserved *cis*-elements that contribute to their adaptability to host environment (Sanchez-Vallet et al., 2018, Dong et al., 2016). Thus, H3K27me3 seems to preferentially modify the poorly conserved or newly evolved *effectors* to “guard” the genome through transcriptional silencing (Dong et al., 2015, Sanchez-Vallet et al., 2018, Fouche et al., 2018). Whenever necessary, such as in response to the host environment, fungal pathogens redistribute H3K27me3 and evoke transcriptional activation, then the activated *effectors* help the pathogens to better adapt to and colonize their host (Fouche et al., 2018).

Polycomb silencing has been extensively studied in multicellular organisms, which is conserved from *Drosophila* to mammals and higher plants, as well as in the fungi kingdom (Margueron and Reinberg, 2011, Ridenour et al., 2020, Wiles and Selker, 2017, Schuettengruber et al., 2017). The core subunits Kmt6, Suz12 and Eed are necessary for H3K27me3 modification and transcriptional silencing (Wiles and Selker, 2017). However, the role of *P55*, the additional subunit of PRC2, remains obscure in fungi. In *F. graminearum*, the homolog of subunit p55 is not involved in H3K27me3 modification (Ridenour et al., 2020). In *N. crassa*, *P55* (*NPF*) is critical for H3K27me3 in a region-dependent manner (Jamieson et al., 2013). In our research, we found that *P55* plays a critical role in H3K27me3-mediated transcriptional silencing in *M. oryzae.* First, P55 acts as the bridge to connect three core components together. Second, disruption of *P55* largely reprograms gene expression, which significantly overlaps with those of Δ*kmt6* and is directly associated with H3K27me3 occupancy. Third, P55 recruits Sin3-HDAC to form the co-suppression complex with PRC2.

The Sin3-HDAC corepressor complex is usually associated with a large number of DNA-binding transcription factors or corepressors to achieve specific and timely regulation of local chromatin and transcription (Grzenda et al., 2009). In Sin3 histone deacetylase complex, Sap30 serves as the bridge and stabilizing molecule and Sap18 also involves in the apoptosis and splicing associated protein (ASAP) complex (Adams et al., 2018, Grzenda et al., 2009, Julia I. Qüesta, 2016). In our study, we found that P55 recruits Sin3 histone deacetylase complex to form co-suppression complex. Furthermore, Sin3 functions as a transcriptional repressor and participates in the *effectors*-related biological process similar to PRC2. All these results suggested that Sin3-HDAC would promote H3K27me3 occupancy and facilitate PRC2 machinery in stably maintaining condensed chromatin status and achieving gene silencing coordinately. In addition, it is possible that Sin3-associated transcription factors or corepressors help H3K27me3 recruitment properly (Adams et al., 2018). This finding not only provides a plausible explanation that PRC2 could stably maintain transcriptional silencing in the absence of PRC1, but also elucidates the molecular clue of transcription output caused by crosstalk between histone methylation and (de)acetylation in *M. oryzae*.

Establishment and maintenance H3K27me3 to local chromatin depends on histone “reader” and series of *trans*/*cis*-regulatory factors which act in corporate or alone (Kassis and Brown, 2013, Xiao et al., 2017). One of the best example of epigenetically reprograming is regulation of floral repressor *Flowering Locus C* (*FLC*) under vernalization (winter cold exposure) in model plant *Arabidopsis* (Luo and He, 2020). In our study, we found that the silenced effectors would be de-repressed with reduced H3K27me3 occupancy during invasive growth stage. Consistent with *FLC* as molecular switch to “remember” vernalization in *Arabidopsis* (Luo and He, 2020), transcriptional reprogramming of some specific *effectors* in *M. oryzae* may act as the molecular switch to “remember” the host environment. However, it is still unknown how the repressed chromatin is “erased” and how the activated state is established during invasive growth stage. Given the destined *effectors*, it is worthy to further elucidate these detailed mechanisms.

## Materials and Methods

### Fungal strains and culture conditions

The *M. oryzae* WT strain B157 and Guy11 were used as background strain to obtain transformants in this study, which were kind gifts from the Indian Institute of Rice Research (Hyderabad, India). All the WT in this study is B157, except for those marked with Guy11. For growth assessment of *M. oryzae,* strains were grown on the prune agar medium (PA) for 7 d (Kou et al., 2017). For conidiation, strains were grown on CM medium at 28℃ in the dark for 2 d, followed by growth under continuous lights for 5 d.

### Plasmid construction

To create the deletion mutants of *KMT6*, *Eed*, *Suz12*, *P55*, *Sin3*, *Sap18* and *Sap30*, the standard one-step gene replacement strategy was used in this study. Briefly, approximate 1-kb of 5’UTR and 3’UTR regions were amplified and ligated sequentially to the flanking of *Hygromycin*, *Sulf* or *Bar* gene cassette in the *pFGL821* (Addgene, 58223), *pFGL820* (Addgene, 58222) or *pFGL822* (Addgene, 58225) respectively (Kou et al., 2017). The sequences of plasmids were confirmed by sequencing and subsequently introduced into the B157 strain by *Agrobacterium tumefaciens* mediated transformation (ATMT). To generate the localization and complementary vectors, the *eGFP* and *TrpC* terminator were cloned to *pFGL820* to obtain *pFGL820*-*GFP-TrpC* terminator. The fragments containing about 1.5-kb of promotor and coding region were amplified (primers listed in Table S2) and then cloned to *pFGL820*-*GFP-TrpC* terminator. After conformed by sequencing, the resultant plasmids were introduced into corresponding deletion mutants respectively by ATMT.

To generate the constructs of *pRP27-KMT6-Flag*, *pRP27-Suz12-Flag* and *pRP27-P55-Flag* for co-immunoprecipitation assay, the coding sequence were cloned to *pFGL822-pRP27-Flag* or *pFGL820-pRP27-eGFP*. The confirmed plasmids were introduced into deletion mutants respectively by ATMT. Primers used in the experiments were listed in Table S2.

### Live cell imaging and image processing

The CM cultivated hypha were stained with 100 μg/mL Hoechst 33342 (Sigma, 14533) for 20 min to visualize nuclei. Live cell epifluorescence microscopy imaging was performed with LSM700 (CarlZeiss Inc.), using the requisite conditions established for detecting GFP or Hoechst signals. Image processing was performed using image J (http://fiji.sc/wiki/index.php/Fiji). These experiments were conducted with two independent repeats.

### Appressorial formation assay

For appressoria formation assay, conidia were harvested from 7-d-old cultures and resuspended in sterile water at a concentration of 10^5^ conidia per mL. 10 μL droplets of the conidial suspension were inoculated on the hydrophobic plastic cover slips and incubated in high humidity at room temperature (Kou et al., 2017). The appressoria were quantified at 24 hpi. Photographs were taken with an Olympus BX53 wide field microscope equipped with bright field optics.

### Rice seedling and rice sheath infection assay

Rice seedling infection assays were performed as described with 5×10^4^/ mL conidial suspension (Kou et al., 2017). Disease symptoms of infection assays were assessed on 7 dpi. The infection assays were repeated three times. For the invasive hyphal development assay, 21-d-old healthy rice seedlings (CO39) were used for sheath preparation. Conidial suspension was inoculated into the rice sheath and incubated in the growth chamber with the photoperiod of 16-h light and 8-h dark at 25°C. The inoculated sheath was trimmed manually and observed by using an Olympus BX53 wide field microscope at 40 hpi.

### Yeast two-hybrid analysis

Yeast two-hybrid assays were performed in the yeast strain Y2Hgold with Matchmaker two-hybrid system (Clontech) according to manufacturer’s instruction. The baits and preys were cloned to pGADT7 and pGBKT7 respectively. The transformants were grown on the basic medium without tryptophan and leucine at 30°C for about 2 d, and subsequently the direct interaction between two proteins were tested with grown on the selective medium without tryptophan, leucine, histidine and adenine for 4-6 d.

### Western blotting analysis

For detecting Flag/GFP-tagged proteins, 0.1 g mycelia cultured in the liquid CM for 2 d were collected. Total protein was extracted with lysis buffer (50 mM Tris-HCl pH 7.4, 150 mM NaCl, 1 mM EDTA, 1% Triton 100 and 1 × protein inhibitor). The resulting protein was separated by 8-15% SDS polyacrylamide gel electrophoresis (SDS-PAGE) and transferred to PVDF membrane, subsequently detected by immunoblotting with anti-Flag (Sigma, A8592) or anti-GFP (Abcam, ab290) antibody.

For detecting histones, the nuclei of 0.5 g mycelia were isolated with extraction buffer (20 mM Tris pH 7.5, 20 mM KCl, 2 mM MgCl_2_, 25% Glycerol, 250 mM Sucrose, 0.1 mM PMSF, 5 mM beta-mercaptoethanol, 1 × proteinase inhibitors) and filtered through two layers of Miracloth (Millipore). Then the histones were extracted with lysis buffer (50 mM Tris-HCl pH 7.4, 150 mM NaCl, 1 mM EDTA, 1% Triton 100 and 1 × protein inhibitor). Total histones were separated by 15% SDS-PAGE gel and protein blots were detected with anti-H3 (Millipore, 06-755), anti-H3K27me3 (Abcam, ab6002), anti-H3K4me3 (Abcam, ab1012), anti-H3K36me3 (Abcam, ab9050), anti-H3K4ac (Active motif, 39381), anti-H3K9ac (PTMBIO, PTM-112), anti-H3K18ac (PTMBIO, PTM-158), anti-H3K27ac (Abcam, ab177178) and anti-H4K5ac (Millipore, 07-327) antibody. The relative intensity of Western blots was quantified by Image J software.

### Co-immunoprecipitation assays

To perform co-immunoprecipitation assays, 0.5 g mycelia cultured in the liquid CM for 2 d were harvested. Total protein was extracted with lysis buffer (50 mM Tris-HCl pH 7.4, 150 mM NaCl, 1 mM EDTA, 1% Triton 100 and 1 × protein inhibitor) and subsequently precipitated with anti-Flag M2 Magnetic beads (Sigma, M8823) or GFP-Trap (Chromotek, gtma-20) respectively according to the manufacture’s instruction. The precipitated protein and input control were detected by the Western blotting with anti-Flag (Sigma, A8592) or anti-GFP (Abcam, ab290) antibody.

### mRNA expression analysis with RT-qPCR

Total RNA was extracted from CM cultivated mycelia for 2 d from three biological repeats using the TRIzol (Invitrogen, USA) reagent. Subsequently total RNAs were reversely transcribed into cDNAs with commercial kits (TOYOBO, FSQ-301) according to the manufacturer’s instruction. RT-qPCR was performed using SYBR Green qPCR Master Mix (TOYOBO, QST-100) in the LightCycler480 system (Roche). The constitutively expressed *tubulin* gene (*MGG_00604*) was used as endogenous control to normalize amount of cDNA templates. Primers used in the experiments were listed and described in Supplemental Table 2.

### RNA-seq analysis

Total RNA was extracted from mycelia from three biological repeats using the TRIzol (Invitrogen, USA) reagent according to the manufacturer’s instruction. RNAs were sequenced separately at Novogene, using Illumina Hiseq X-Ten with Hiseq-PE150 strategy. The clean reads were mapped to the reference genome of *M. oryzae* (*M. oryzae* 70-15 assembly MG8 from NCBI) using TopHat software with default settings (Daehwan Kim, 2013). For analysis of differential expression, the assembled transcripts from three independent biological replicates in the WT and deletion mutants were included and compared using Cuffdiff with default settings (Trapnell et al., 2010). Gene with false discovery rate (FDR) threshold of 0.05 in conjunction with at least 2-fold change in expression level was considered differentially expressed. To calculate the significance of the overlap of two gene sets, P value with Fisher’s exact test for overlapping was performed on online (the total number of genes in the *M. oryzae* genome used was 14317) (http://nemates.org/MA/progs/overlap_stats.html). GO analysis for enriched biological processes was perform using DAVID (https://david.ncifcrf.gov/home.jsp) with default settings.

### ChIP and ChIP-seq analysis

The ChIP experiments with mycelia were conducted as previous reports with minor modification (He et al., 2018, Tao et al., 2017). Briefly, 1 g mycelia were crosslinked with 1% formaldehyde for 20 mins and stopped with 125 mM glycine for 5 mins at room temperature. Samples were ground with liquid nitrogen and resuspended in the nuclei isolating buffer (10 mM Tris pH 8.0, 10 mM Sodium butyrate, 400 mM Sucrose, 0.1 mM PMSF, 5 mM Beta-Mercaptoethanol, 1 × proteinase inhibitors). Subsequently the precipitated nuclei were used to total chromatin extraction with 1 mL lysis buffer (50 mM HEPES pH7.5, 150 mM NaCl, 1mM EDTA, 10 mM Sodium butyrate, 0.1% deoxycholate, 0.1% SDS, 1% Triton X-100, 1mM PMSF and 1 × Roche protease inhibitor Cocktail). The obtained chromatin was sonicated into DNA fragments between 200-500 bp using Diagenode Bioruptor (high setting, 16 cycles, every cycle with 30 seconds “on” and 30 seconds “off”). 20 μL chromatin solution? was used to input DNA extraction and the remainder was pre-cleared with 10 μL protein A Dynabeads (Thermofisher, 10001D) for 1 h. Then, the chromatin was incubated with anti-H3K27me3 (Abcam, ab6002) or anti-H3K4ac (Active Motif, 39381) overnight at 4°C. Another 20 μL protein A Dynabeads was used to capture protein-DNA mixture and washed for three times. Protein-DNA mixture was reverse-crosslinked, and DNA was recovered with phenol-chloroform extraction. The recovered DNA was used as template for followed ChIP-qPCR and ChIP-seq. Two biological repeats were carried out.

For ChIP-seq assay, the purified DNA was used as library construction with the NEBNext Ultra II DNA Library Prep Kit for Illumina (NEB, E7645L). High-throughput sequencing was carried out using Illumina Hiseq-PE150 by Novogene Corporation (Beijing, China) for Illumina (Langmead et al., 2009). Subsequently the clean read pairs were mapped to the reference genome with Bowtie2 (Version 2.2.8) (Langmead and Salzberg, 2012) and enriched peaks were called by MACS2 (Version 2.1.1) with default parameters (Zhang et al., 2008). The data was imported into the integrative genomics viewer (IGV) for visualization (James T Robinson, 2011). To assign peaks to proximal genes, the distance of 3-kb flanking the peak summit were extracted. To validate ChIP-seq results, ChIP-qPCR assay with two independent repeats was performed. The level of examined fragments was relative to internal reference gene *TUB5* using quantitative real-time PCR. The PCR primers were listed and described in Table S2.

### Data availability

The ChIP-seq and RNA-seq datasets generated in this article were deposited in the Gene Expression Omnibus (GEO) under the accession number GSE166690.

## Acknowledgments

We thank Dr. Naweed I. Naqvi of the National University of Singapore for helpful suggestions and critically reading manuscript. Thanks to Dr. Zhou Ming of Zhejiang University for help of bioinformatics analysis. Thanks to Dr. Qi Zhenyu of agricultural experimental station of Zhejiang University for controlling the light environment of artificial climate chamber.

## Author contributions

Y.K. and Z.T. conceived and designed the experiments; Y.K., J.Q. and Z.W. performed the experiments; H.S., C.L., J.Y., Z.L. and W.X. provided technical assistance and contributed reagents/materials/analysis tools; Y.K. and Z.T. wrote the paper.

## Supporting Information

The following Supporting Information is available for this article:

**Figure S1.**
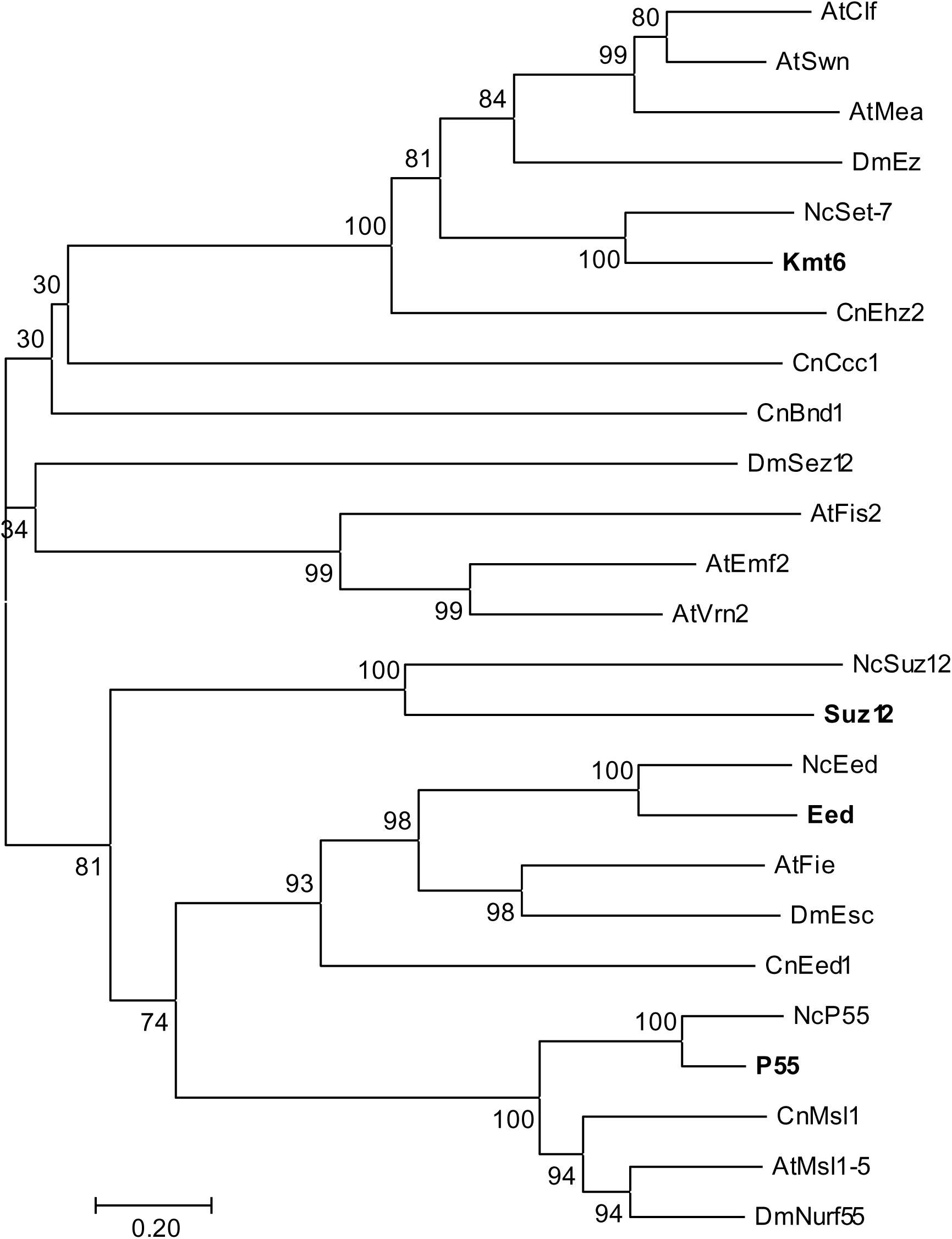
Phylogenetic analysis of PRC2 components in *M. oryzae*. The phylogenetic tree of PRC2 components from *M. oryzae* (highlighted with bold font), *N. crassa*, *A. thaliana*, and *D. melanogaster* was constructed by CLUSTAL_W and MEGA 7 programs using neighbor-joining method with 1000 bootstrap replicates. Genbank accession numbers of these protein were listed in S1 Table.

**Figure S2.**
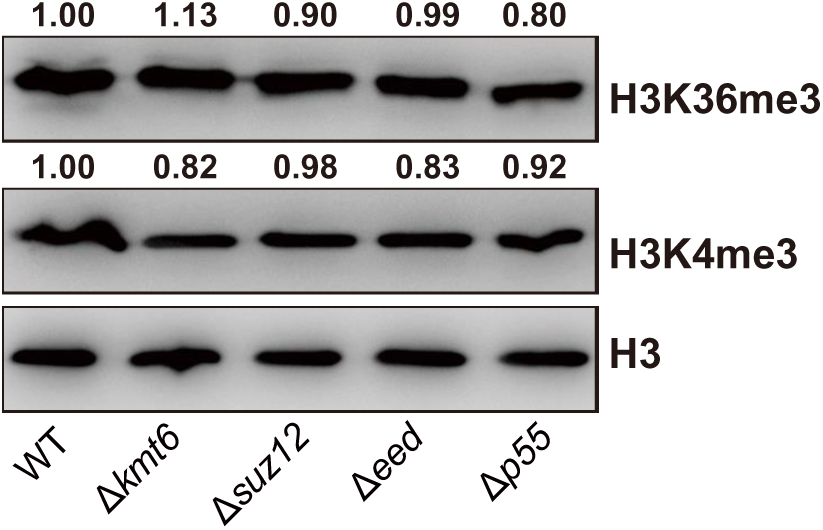
Detection of H3K4me3 and H3K36me3 level in the deletion strains by Western blot analysis. The number at the top of band was the relative intensity calculated with Image J software. Two biological replicates were carried out with similar results.

**Figure S3.**
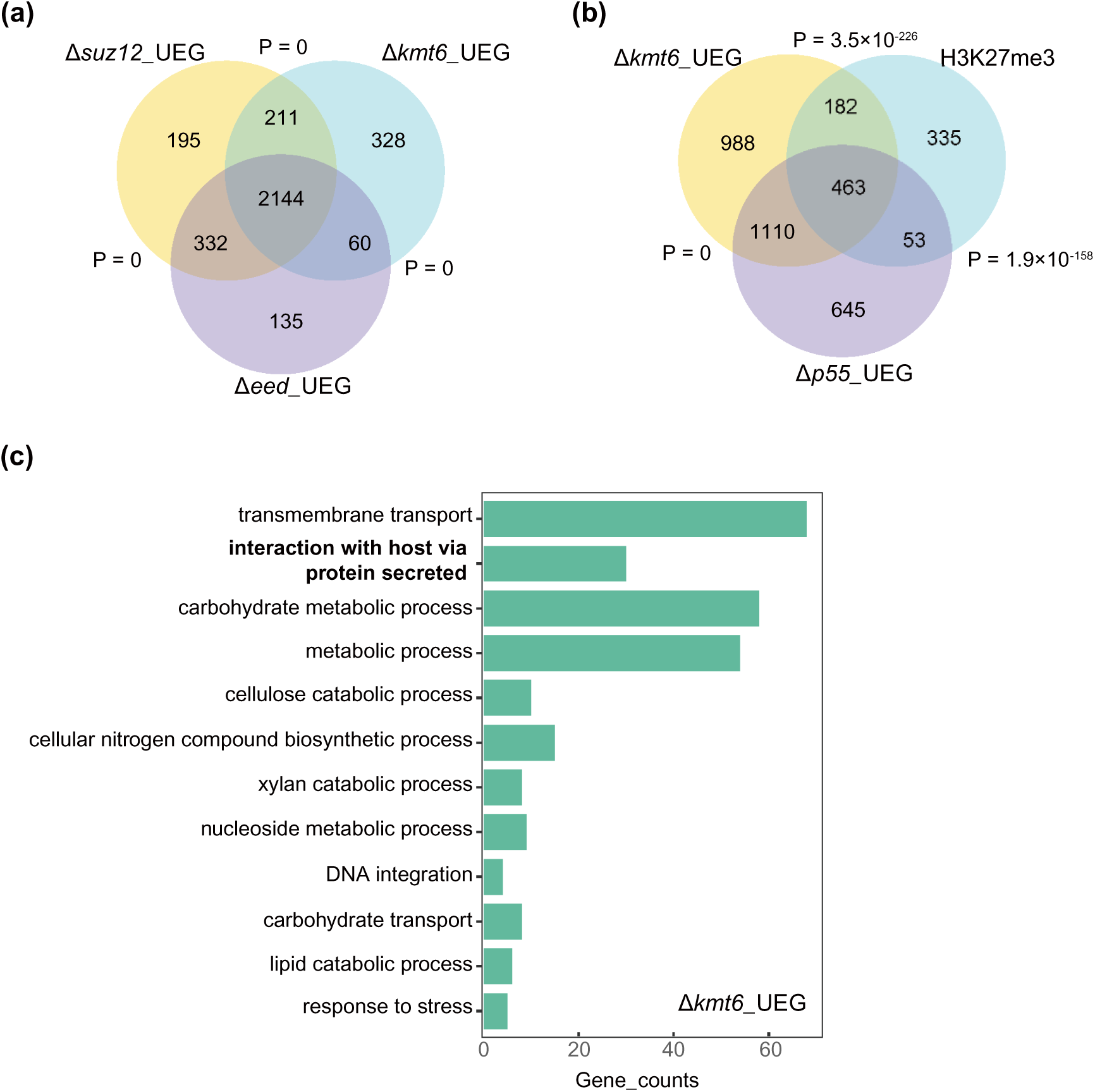
Bioinformatics analysis of PRC2 subunits based on RNA-seq. (a) The Venn diagram showing statistically significant overlaps among genes sets of the Δ*kmt6*-UEG, Δ*suz12*-UEG and Δ*eed*-UEG. P value for overlapping between gene sets was obtained by Fisher’s exact test. (b) The Venn diagram showing significant overlap among gene sets of Δ*kmt6*-UEG, Δ*p55*-UEG and H3K27me3 marked genes. (c) GO analysis on gene sets of the Δ*kmt6*-UEG.

**Figure S4.**
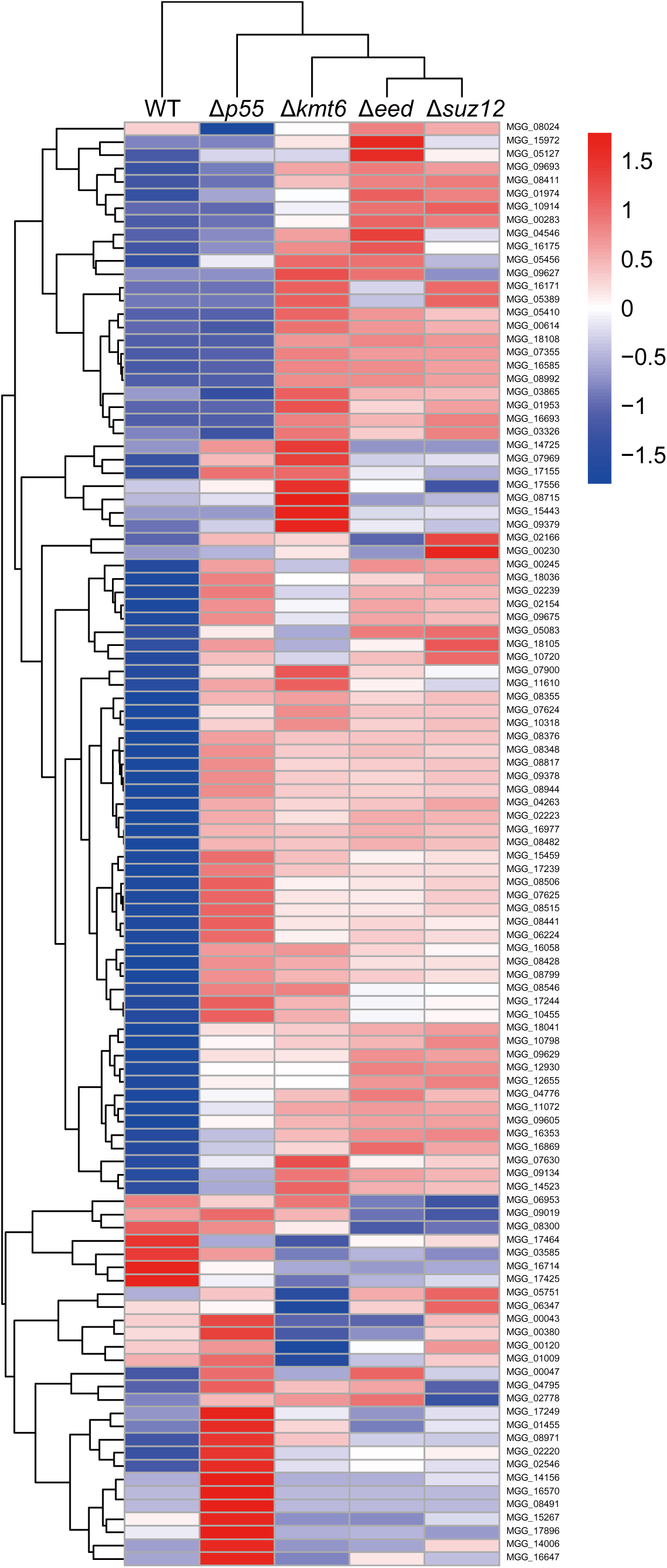
Expression pattern of *effectors* in the *PRC2* mutants. Heat maps illustrated expression changes of genome-wide e*ffectors* in different *PRC2* mutant compared with that of WT B157. Three biological replications of RNA-Seq sequencing were conducted on independent samples.

**Figure S5.**
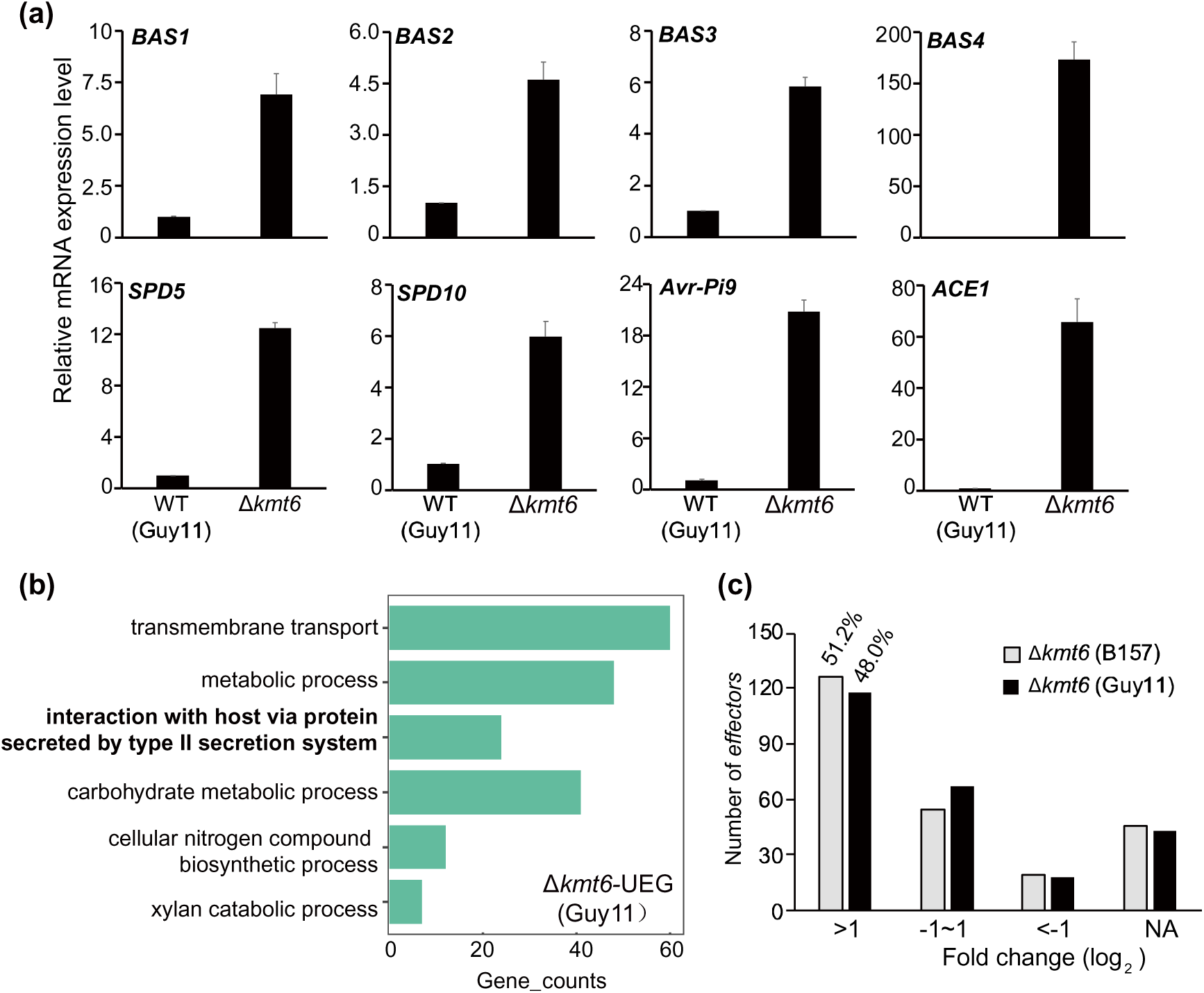
Expression pattern of *effectors* in the Δ*kmt6*(Guy11) mutant. (a) Relative expression level of representative *effectors* in Guy11 and Δ*kmt6*(Guy11). Values are means ± SD of three biological replicates. (b) GO analysis on gene sets of the Δ*kmt6*-UEG (Guy11). (c) The distribution of putative 246 *effectors* based on the expression level change in the Δ*kmt6*-UEG (B157) and Δ*kmt6*-UEG (Guy11).

**Figure S6.**
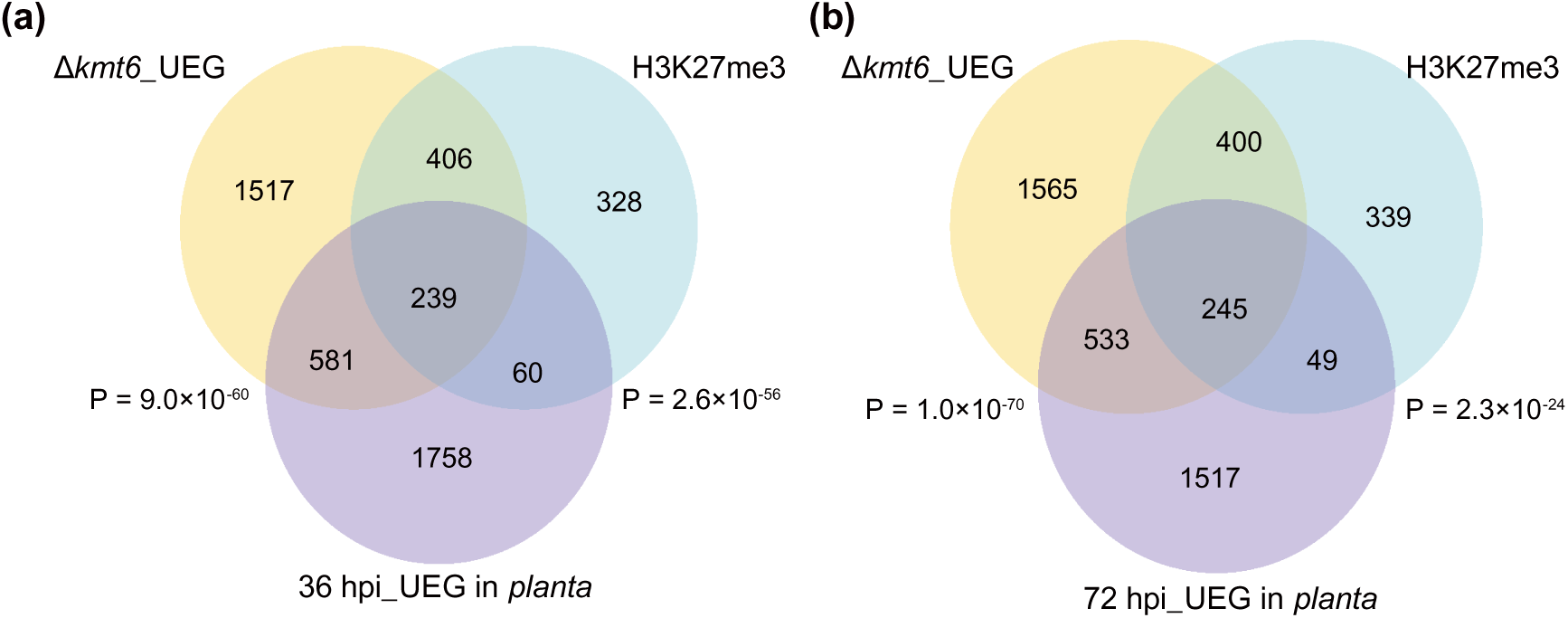
Absence of H3K27me3 partially mimics host-derived signal. (a) The Venn diagram shows statistically significant overlaps among genes sets of the Δ*kmt6*-UEG, 36 hpi_UEG *in planta* and H3K27me3 marked genes. (b) The Venn diagram represents statistically significant overlaps among genes sets of Δ*kmt6*-UEG, 72 hpi_UEG *in planta* and H3K27me3 marked genes.

**Figure S7.**
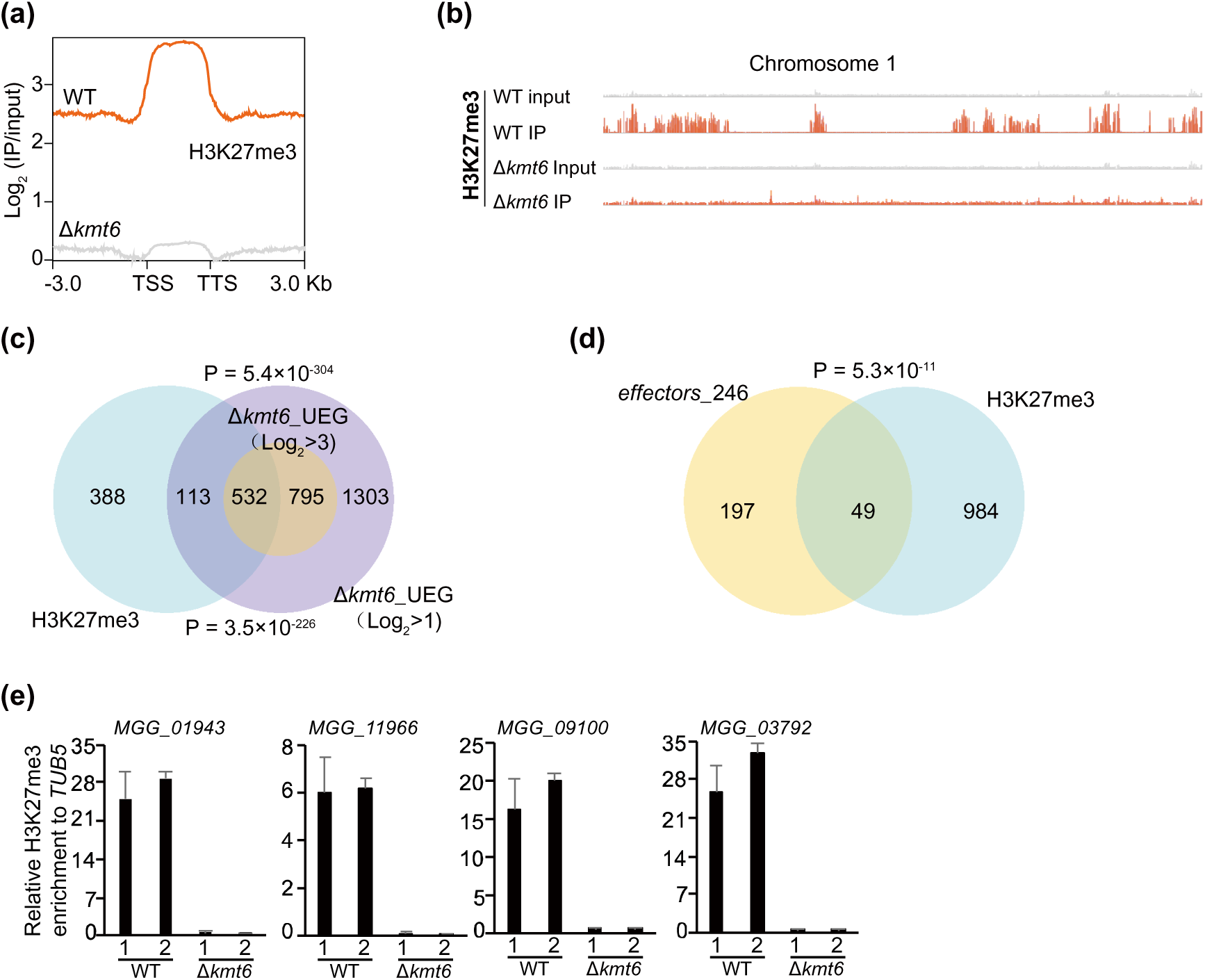
H3K27me3 modification deposits on the chromatin of *effectors* and infection-specific genes during vegetative growth stage. (a) The average H3K27me3 occupancy within 3 Kb genomic regions flanking the summit of H3K27me3 peaks in the WT and Δ*kmt6* strains. (b) Genome browser views of H3K27me3 occupancy on the chromosome 1 in the WT and Δ*kmt6* strains. (c) The Venn diagram shows statistically significant overlaps between gene sets of H3K27me3 marked genes and the Δ*kmt6*_UEG with threshold of Log_2_ > 1 and Log_2_ > 3 respectively. P value with Fisher’s exact test for overlapping between gene sets were labeled. (d) The Venn diagram shows significant overlap among gene sets of putative 246 *effectors* and H3K27me3 marked genes. (e) ChIP-qPCR verified the enrichment of H3K27me3 on the chromatin of infection-specific genes. The relative enrichments were calculated by relative quantitation of two biological repeats, which was standardized by internal control *TUB5*, and then compared with that of WT. Bar represents standard error from three technical repeats.

**Figure S8.**
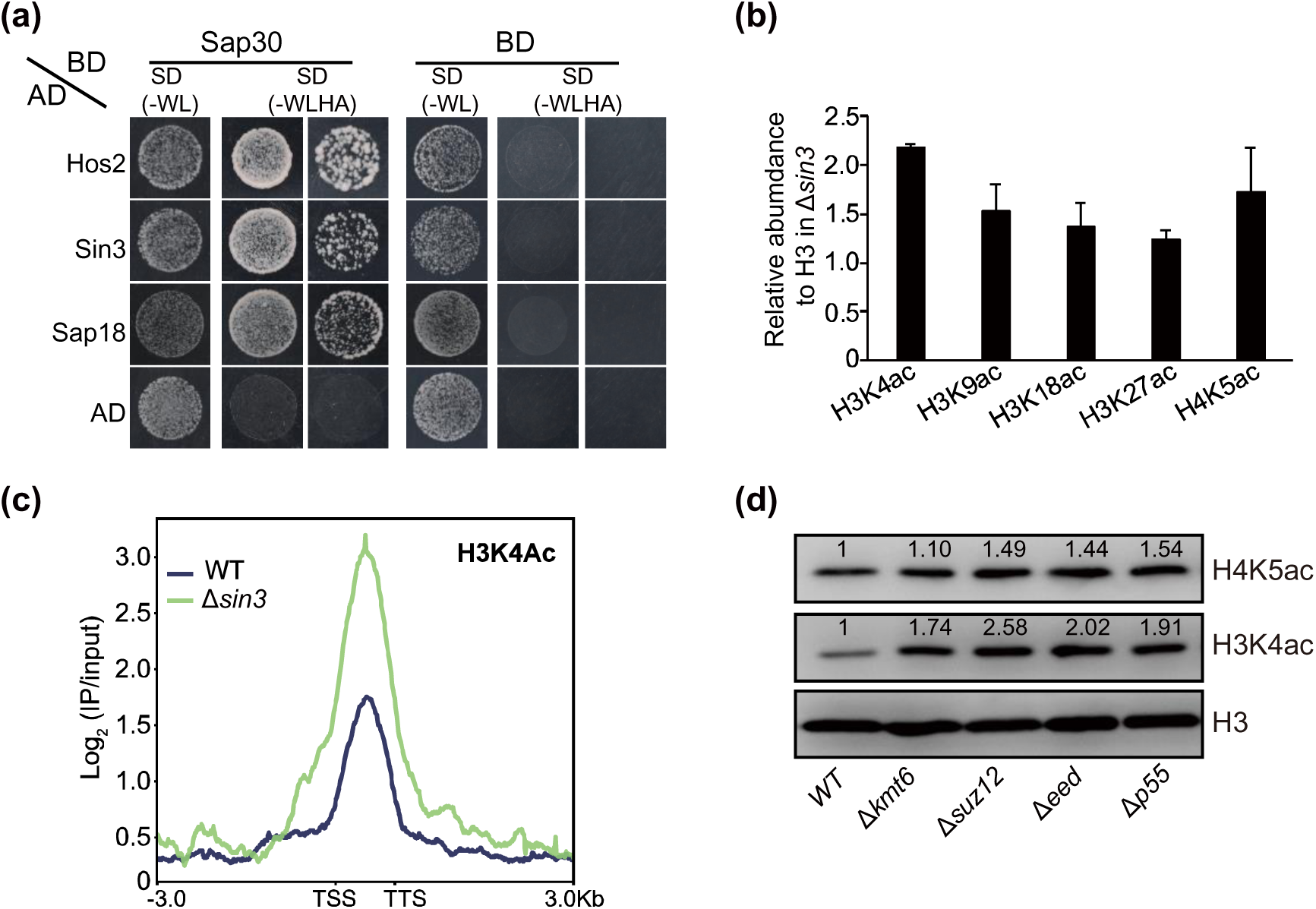
Sin3 histone deacetylase complex contributes to histone deacetylation. (a) Yeast-two-hybrid assay of interaction between Sap30 and other Sin3-HDAC components Hos2, Sin3, or Sap18. The bait and prey plasmids were co-transformed into yeast strain Y2Hgold respectively, then the transformants were grown on basal medium SD (-WL) and selective medium SD (-WLHA). (b) Relative abundance of examined protein blots in the WT and Δ*sin3* strains. The standard error was obtained from three biological repetitions. (c) Bolt representation of the average H3K4ac occupancy within 3 Kb genomic regions flanking the summit of H3K4ac peaks in the WT and Δ*sin3* mutant. (d) Western blotting assay detected the H4K5ac and H3K4ac modification in the deletion strains of PRC2 subunits. The number at the top of band was the relative intensity calculated with Image J. Two biological replicates were carried out with similar results.

**Figure S9.**
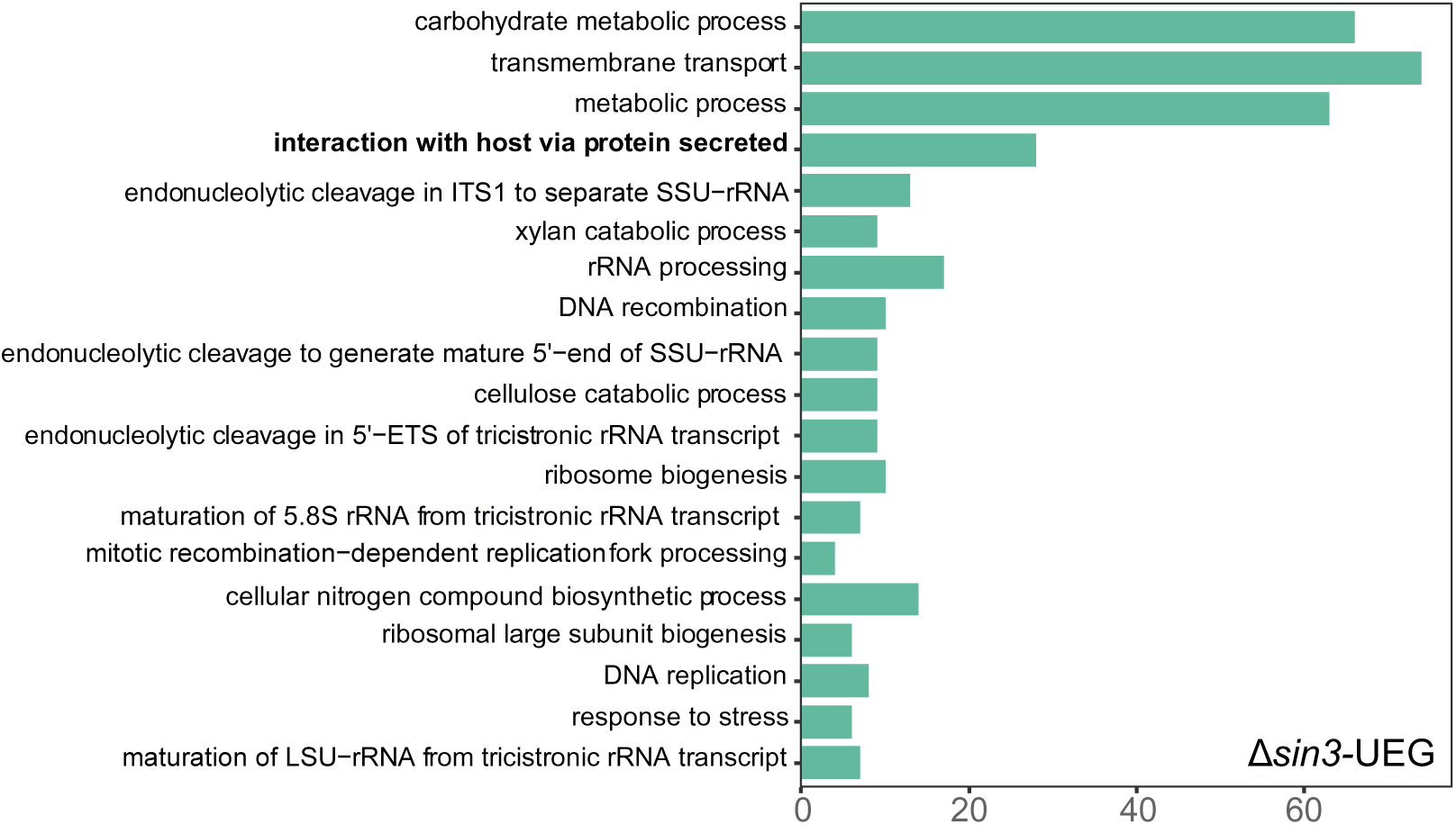
GO analysis on gene sets of Δ*sin3*-UEG.

**Figure S10.**
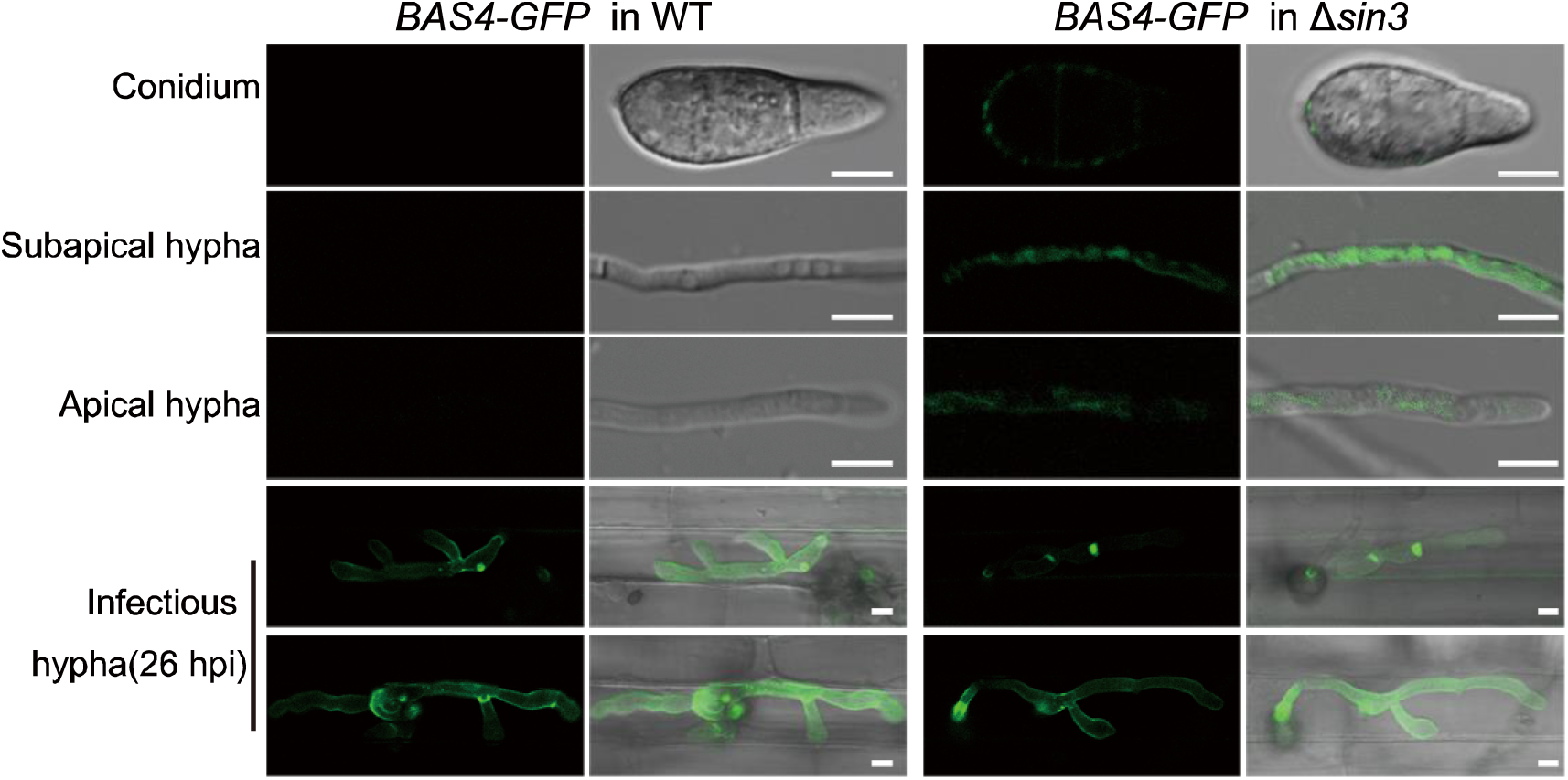
Confocal microscopy-based subcellular localization of Bas4-GFP in the vegetative and *in planta* growth stages in the WT and Δ*sin3* strains. In the WT, Bas4-GFP fluorescence was only presented in the *in planta* growth stage, while in the Δ*sin3*, Bas4-GFP fluorescence could be detected during both vegetative and *in planta* growth stages, including the conidia, apical hypha, subapical hypha and invasive hypha. Bar = 5 μm.

**Table S1.**
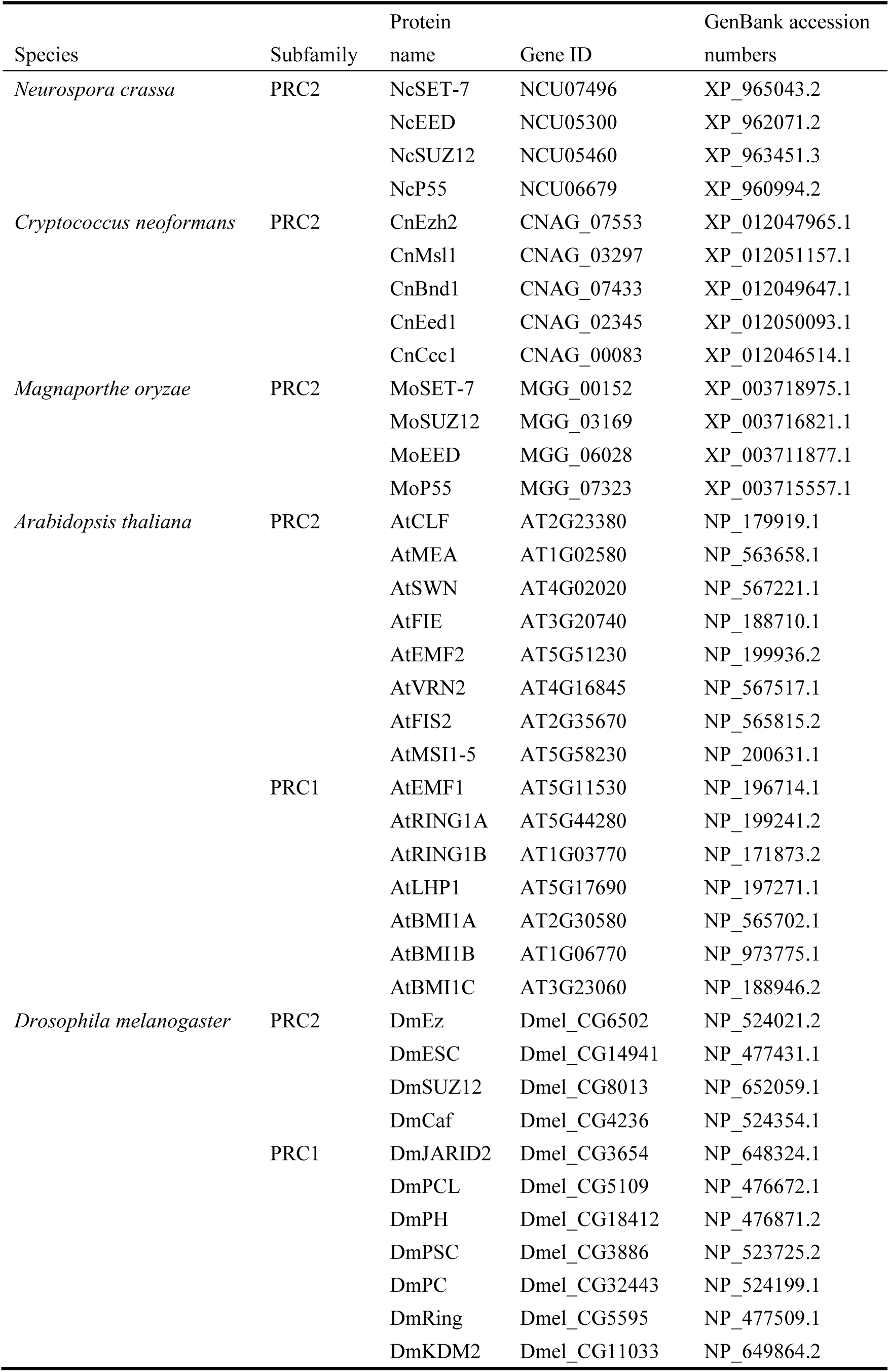
Components of polycomb repressive complex in *M. oryzae*, *N. crassa*, *A. thaliana*, and *D. melanogaster*.

**Table S2.**
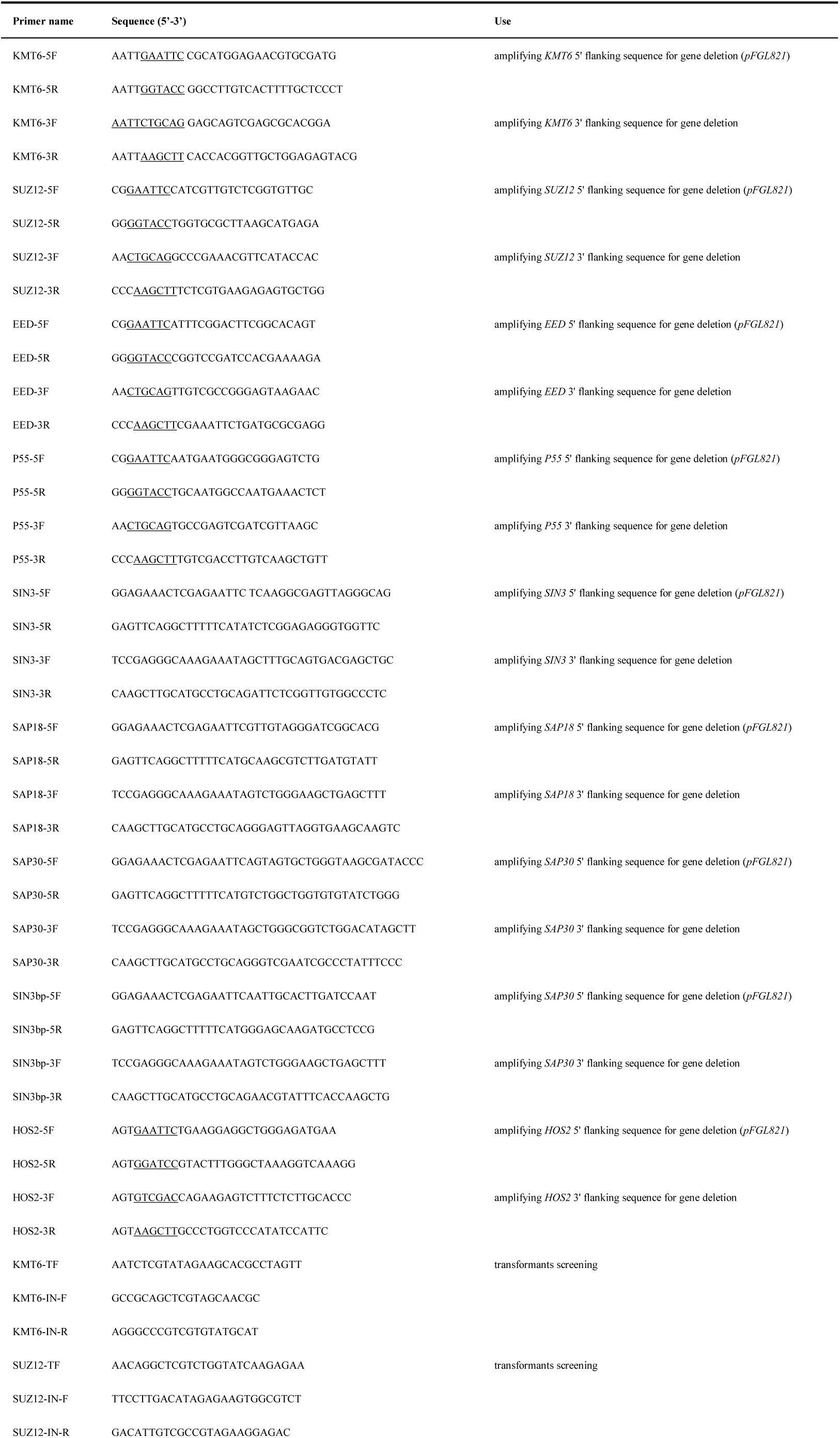
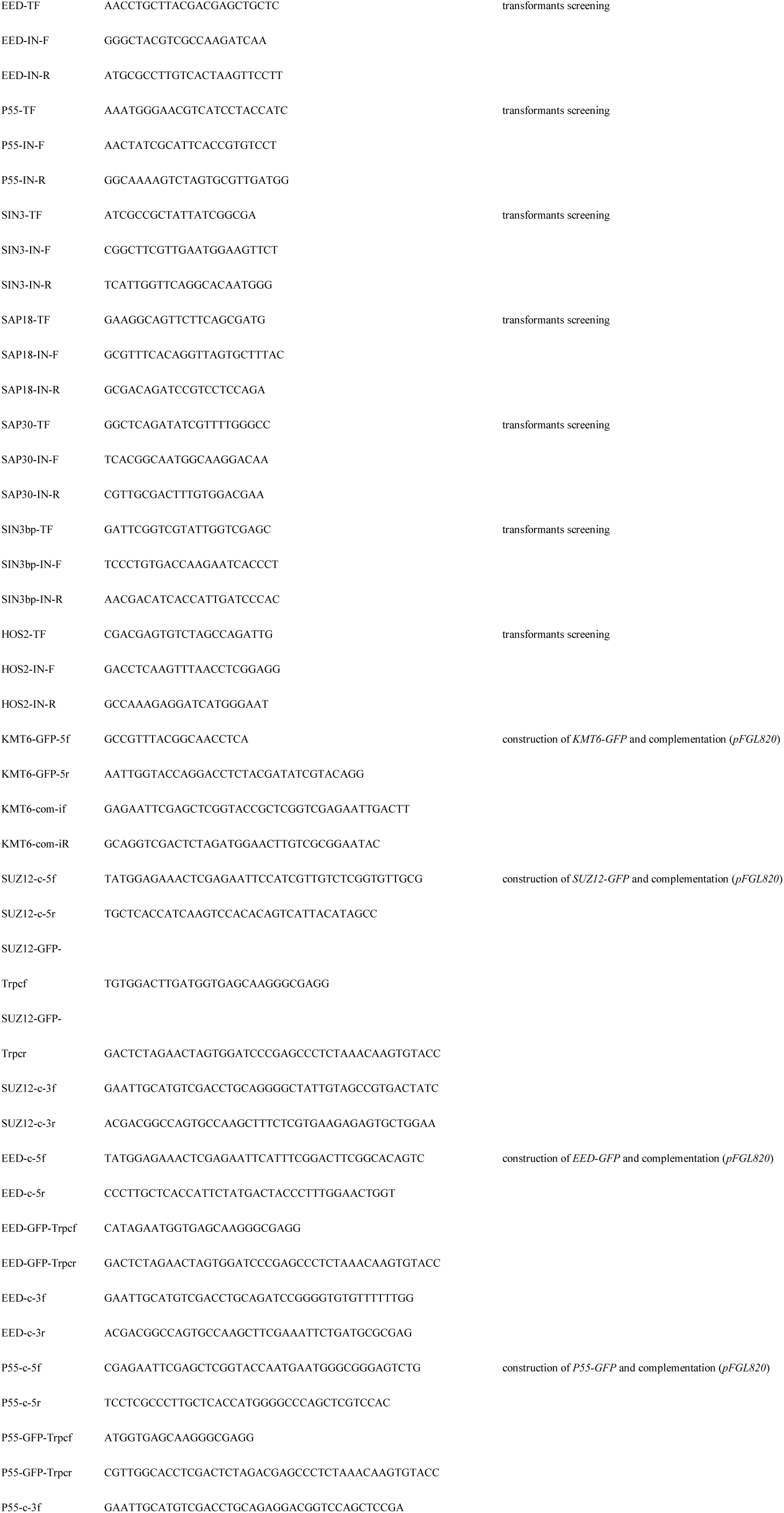
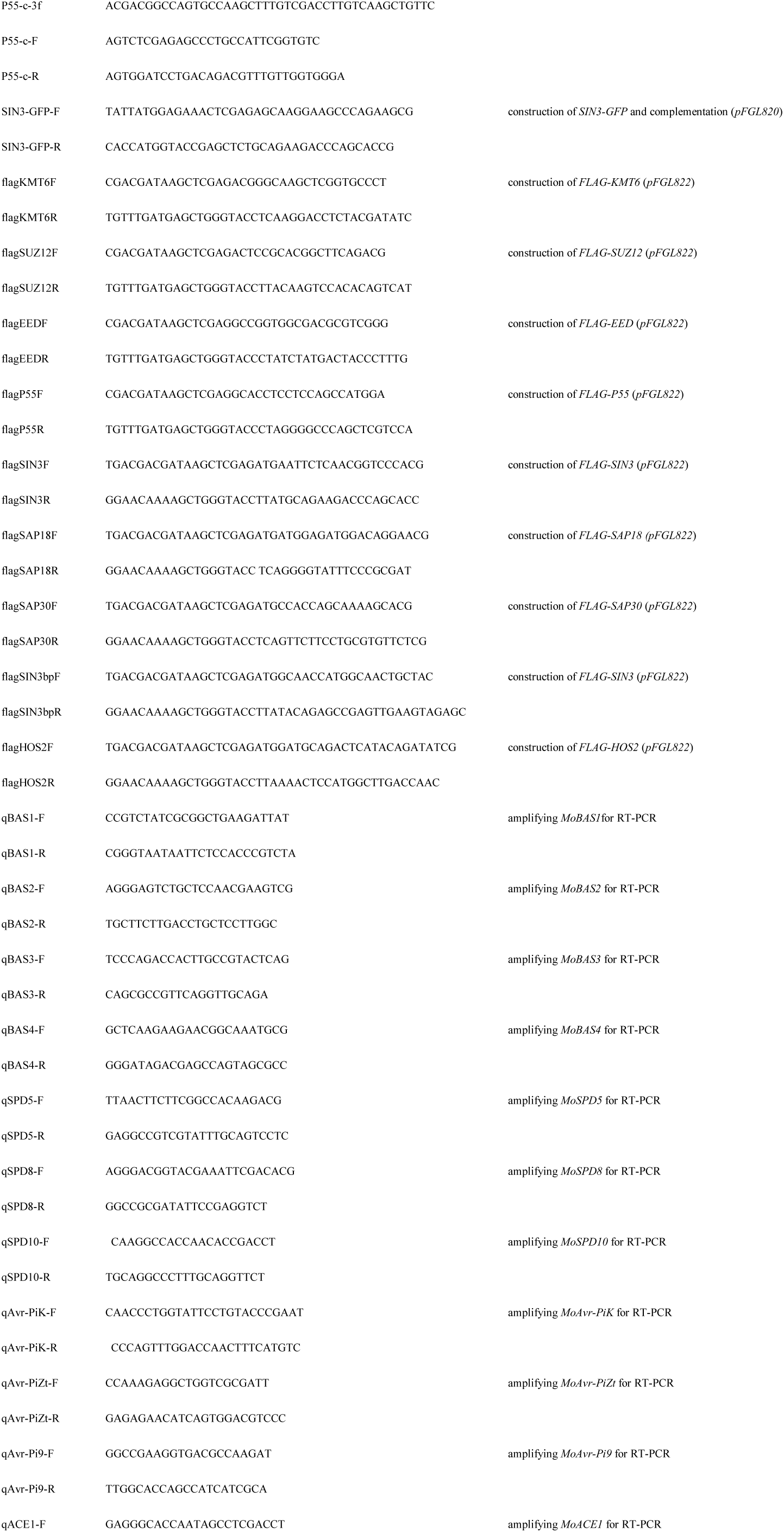
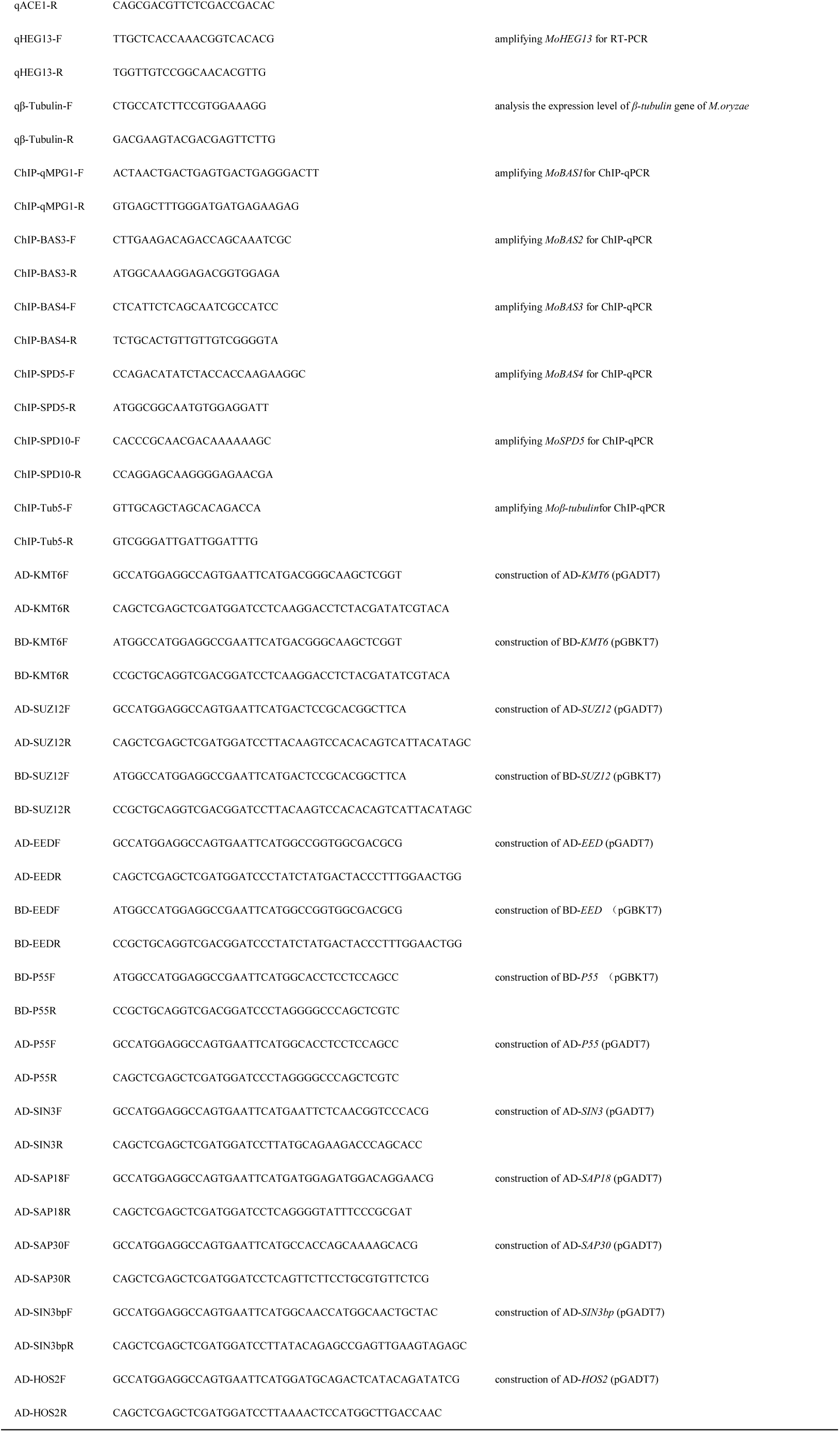
Primers used in this study.

**Table S3.**
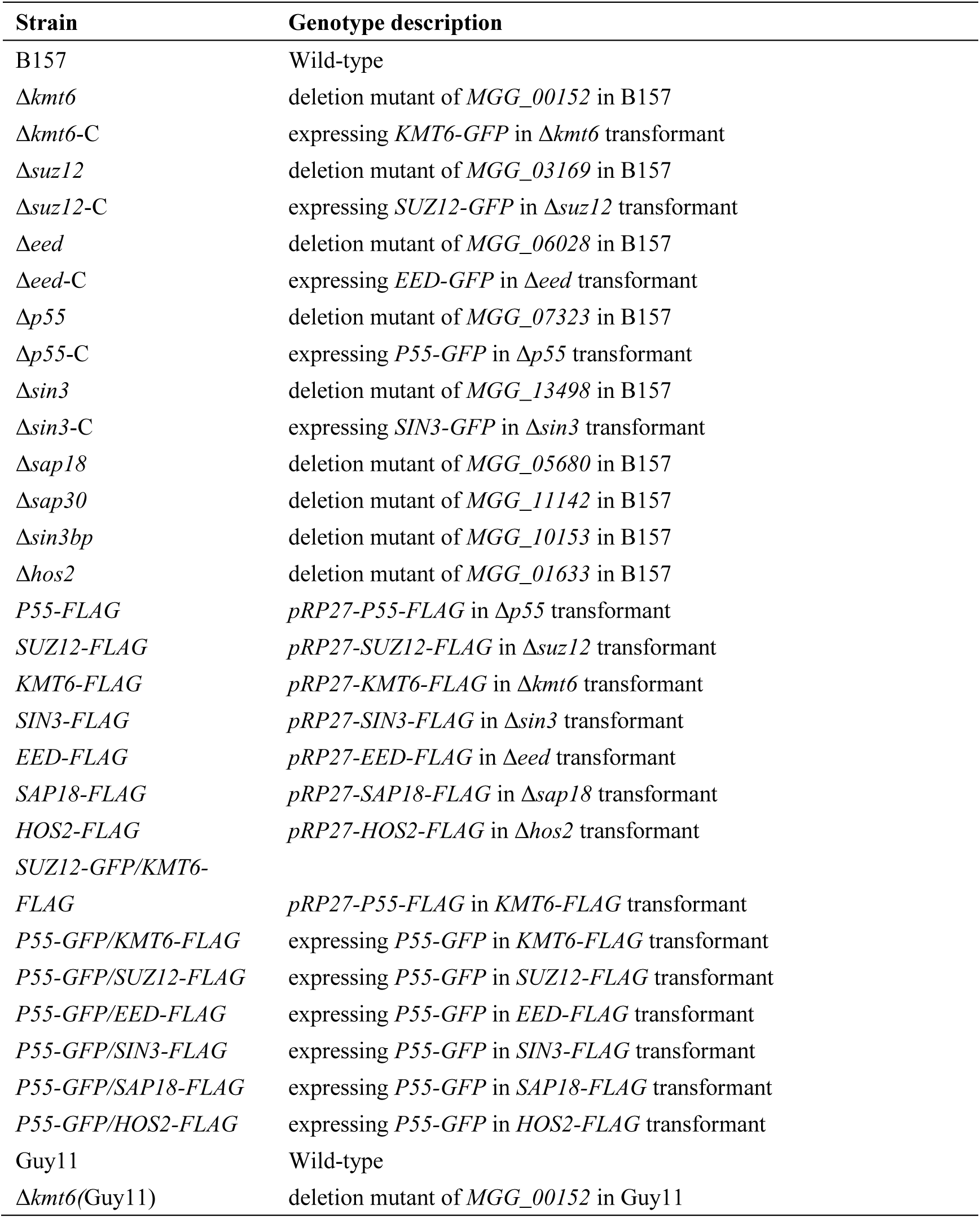
Strains used in this study.

| Strain | Genotype description |
| --- | --- |
| B157 | Wild-type |
| $\Delta kmt6$ | deletion mutant of *MGG_00152* in B157 |
| $\Delta kmt6$-C | expressing *KMT6-GFP* in $\Delta kmt6$ transformant |
| $\Delta suz12$ | deletion mutant of *MGG_03169* in B157 |
| $\Delta suz12$-C | expressing *SUZ12-GFP* in $\Delta suz12$ transformant |
| $\Delta eed$ | deletion mutant of *MGG_06028* in B157 |
| $\Delta eed$-C | expressing *EED-GFP* in $\Delta eed$ transformant |
| $\Delta p55$ | deletion mutant of *MGG_07323* in B157 |
| $\Delta p55$-C | expressing *P55-GFP* in $\Delta p55$ transformant |
| $\Delta sin3$ | deletion mutant of *MGG_13498* in B157 |
| $\Delta sin3$-C | expressing *SIN3-GFP* in $\Delta sin3$ transformant |
| $\Delta sap18$ | deletion mutant of *MGG_05680* in B157 |
| $\Delta sap30$ | deletion mutant of *MGG_11142* in B157 |
| $\Delta sin3bp$ | deletion mutant of *MGG_10153* in B157 |
| $\Delta hos2$ | deletion mutant of *MGG_01633* in B157 |
| *P55-FLAG* | *pRP27-P55-FLAG* in $\Delta p55$ transformant |
| *SUZ12-FLAG* | *pRP27-SUZ12-FLAG* in $\Delta suz12$ transformant |
| *KMT6-FLAG* | *pRP27-KMT6-FLAG* in $\Delta kmt6$ transformant |
| *SIN3-FLAG* | *pRP27-SIN3-FLAG* in $\Delta sin3$ transformant |
| *EED-FLAG* | *pRP27-EED-FLAG* in $\Delta eed$ transformant |
| *SAP18-FLAG* | *pRP27-SAP18-FLAG* in $\Delta sap18$ transformant |
| *HOS2-FLAG* | *pRP27-HOS2-FLAG* in $\Delta hos2$ transformant |
| *SUZ12-GFP/KMT6-FLAG* | *pRP27-P55-FLAG* in *KMT6-FLAG* transformant |
| *P55-GFP/KMT6-FLAG* | expressing *P55-GFP* in *KMT6-FLAG* transformant |
| *P55-GFP/SUZ12-FLAG* | expressing *P55-GFP* in *SUZ12-FLAG* transformant |
| *P55-GFP/EED-FLAG* | expressing *P55-GFP* in *EED-FLAG* transformant |
| *P55-GFP/SIN3-FLAG* | expressing *P55-GFP* in *SIN3-FLAG* transformant |
| *P55-GFP/SAP18-FLAG* | expressing *P55-GFP* in *SAP18-FLAG* transformant |
| *P55-GFP/HOS2-FLAG* | expressing *P55-GFP* in *HOS2-FLAG* transformant |
| Guy11 | Wild-type |
| $\Delta kmt6$(Guy11) | deletion mutant of *MGG_00152* in Guy11 |

